# Retinoic acid and androgens interact to regulate spermatogenesis in a non-mammalian vertebrate lacking *stra8*

**DOI:** 10.1101/462408

**Authors:** Diego Crespo, Luiz H.C. Assis, Henk J.G. van de Kant, Sjors de Waard, Diego Safian, Moline S. Lemos, Jan Bogerd, Rüdiger W. Schulz

## Abstract

In mammals, retinoic acid (RA) signaling is critical for spermatogonial differentiation and for entering meiosis, the latter depending on RA-induced *Stra8* gene expression. Many fish species, including zebrafish, do not contain a *stra8* gene, but RA signaling nevertheless is important for sperm production. However, it is not known which stages of spermatogenesis respond to RA. Here, we show in zebrafish that RA promotes spermatogonial differentiation and reduces the apoptotic loss of spermatids, but is not required for meiosis. Some of the RA effects are mediated by other genes, in particular *rec8a*. Surprisingly, androgens can partially compensate for the loss of RA signaling, and we identify a link between the endocrine system and RA signaling: follicle-stimulating hormone (Fsh) stimulates testicular RA production. While RA signaling is relevant at the basis of the vertebrates, it also targets processes and mechanisms that are different from those known in mammals so far.

## Introduction

The production of spermatozoa from spermatogonial stem cells (SSCs) is a developmental process that is regulated by a complex network of endocrine and local signaling pathways in the vertebrate testis. One of the local signaling systems makes use of retinoic acid (RA), the biologically active metabolite of vitamin A, which is crucial for mammalian spermatogenesis at a number of steps (reviewed by (1)). In mice, RA is first required for the transition of undifferentiated to differentiating spermatogonia (2), and again during meiotic initiation, the latter reflecting RA-triggered *Stra8* (stimulated by retinoic acid 8) gene expression in the testis (3, 4). Moreover, STRA8-independent pathways fine-tune RA action in the testis as shown for SALL4A (spalt like transcription factor 4; (5)) and REC8 (REC8 meiotic recombination protein; (6)). A vitamin A (VAD) deficient diet arrested spermatogonial differentiation (7-9), which was reversed by retinoid supplementation (10, 11). Also, mice lacking different isoforms (α, β, γ) of the nuclear RA and retinoid X receptors (RARs and RXRs) showed testicular abnormalities or sterility (12) and germ cell apoptosis (13, 14). Dominant-negative mutant models of the RARα allowed further characterization of essential molecular functions of RA signaling for germ cell development (15, 16). It is not clear if the RA dependency of mammalian spermatogenesis represents an evolutionary conserved feature, as in this regard very little is known in non-mammalian vertebrates including fish, in this regard. Interestingly, *stra8*-like genes are absent in the genomes of a number of model species (including zebrafish [*Danio rerio*] and medaka [*Oryzias latipes*]), so that this RA-responsive factor was suggested to be lost during the evolution of teleost fish (17). More recently, however, *stra8* genes were identified in other fishes. At present, it seems that *stra8* was lost specifically in the Acanthomorpha and Cypriniformes lineages (18). While spermatogenesis proceeds in the absence of *stra8* in these species, in the Southern catfish (*Silurus meridionalis*), for example, entry into meiosis is a *stra8*-sensitive but not a *stra8*-dependent process (19, 20). Intriguingly, in species lacking *stra8*, such as Nile tilapia (*Oreochromis niloticus*) and medaka, RA nevertheless promoted meiotic initiation (21, 22). In zebrafish, RA-producing (*aldh1a2*) and metabolizing (*cyp26a1*) enzymes are expressed in the adult testis (17), and inhibition of RA production by exposure to WIN 18,446 reduced sperm production and fecundity (23), but the males remained fertile. So far, no information has been published regarding the potential regulation of the RA signaling system by reproductive hormones in vertebrates. Also, it is not known what stages of spermatogenesis are sensitive to RA signaling in fish.

In addition to RA signaling, other signaling systems contribute to the regulation of spermatogenesis. Androgens, produced by Leydig cells, are crucial for mammalian spermatogenesis (24, 25) and can initiate or promote germ cell development in GnRH (gonadotropin-releasing hormone)-deficient mice (26) and hypophysectomized rats (27). Furthermore, genetic studies in mice demonstrated that a Sertoli cell-specific androgen receptor (*Ar*) knockout led to meiotic arrest, germ cell apoptosis and the absence of spermatids (28). Meiosis and spermiogenesis (i.e. spermatocytes/spermatids) are particularly dependent on androgen action (reviewed in (29)). Testosterone also promoted spermatogonial differentiation by suppressing the production of WNT5A by Sertoli cells, which would otherwise stimulate SSC self-renewal (30). On the other hand, testosterone stimulated myoid cells to produce GDNF, in turn promoting SSC self-renewal (31). Hence, androgen signaling is also important for the initial, mitotic phase of mammalian spermatogenesis.

In teleost fish, 11-ketotestosterone (11-KT) is considered to be the main androgen (32) and its stimulatory effect on spermatogenesis has been shown in various species such as Japanese eel (*Anguilla japonica*; (33)), Atlantic salmon (*Salmo salar*; (34)), African catfish (*Clarias gariepinus*; (35)) or zebrafish (36, 37). Androgens stimulate the onset of puberty in fish (38, 39) and modulate the testicular expression of genes potentially involved in spermatogenesis (40-42). However, spermatogenesis in fish does not depend as strictly on androgens as in mammals. Loss of the enzyme required for androgen production does not affect spermatogenesis in medaka or zebrafish (43, 44), and loss of *ar* gene function still allows the production of some sperm, although testis weight and sperm numbers become clearly reduced (45, 46). Apparently, pathways operate in the fish testis that support spermatogenesis but do not depend on androgens. In fish, including zebrafish, Fsh is a potent steroidogenic hormone (47) and promotes spermatogonial differentiation via androgens but also via modulating growth factor production by testicular somatic cells (48-53). Hence, androgens, growth factors and RA regulate spermatogenesis, but no information is available on potential interactions between these signaling systems, or whether other reproductive hormones affect RA production. Also, it is not clear how RA signaling is mediated in zebrafish, since a gene of critical relevance for entering meiosis in other vertebrates, *stra8*, is missing in this species. Finally, little information is available about the role of RA effects on zebrafish spermatogenesis.

In the present study, we first carried out mRNA transcriptome profiling to reveal testicular genes/pathways operating during the (re-)start of spermatogenesis following a cytotoxic insult (54). These analyses identified genes of the steroid and RA signaling systems as significantly enriched in this situation. Using primary tissue and cell culture approaches we confirmed retinoid involvement in spermatogenesis, in particular RA-promoted proliferation of differentiating spermatogonia and initiation of meiosis in preleptotene spermatocytes. Genetic evidence suggests that, in the absence of *stra8*, *rec8a*-dependent mechanisms mediate RA action in zebrafish germ cells. While loss of RA signaling in germ cells clearly affects testis histology in 6 month-old transgenic males, it is not critical for sperm production and spermatogenesis recovers at 9 months of age; however, the sperm that are produced are of low quality. Moreover, our studies revealed interactions of RA-signaling and Leydig cells in promoting spermatogenesis. Finally, we found that Fsh stimulates RA production, linking for the first time in vertebrates endocrine regulation of spermatogenesis to RA signaling. Taken together, while our data suggest in general a conservation of the role of RA in stimulating spermatogenesis, in different vertebrate clades different mechanisms are used to implement RA effects.

## Results

### Gene expression profiling during spermatogenic recovery from busulfan treatment

To investigate initial stages of spermatogenesis, this process was first interrupted using the cytostatic agent busulfan in combination with elevated water temperature, and spermatogenesis was then allowed to recover spontaneously from surviving stem cells (54) (*Supplementary file 1A* and *B*). Morphological analysis of testis tissue revealed that maximum germ cell depletion occurred 10 days post-injection (dpi; *Supplementary file 1Biii*). Most spermatogenic tubuli only contained Sertoli cells and very few type A_und_ spermatogonia remained, scattered throughout the testis (*Supplementary file 1Biii* inset). Already 4 days later (i.e. 14 dpi), many tubules contained type A_diff_ and B spermatogonia and spermatocytes (*Supplementary file 1Biv*). This repopulation of the seminiferous epithelium shows the regenerative capacity of the surviving type A_und_ spermatogonia. In recovering testes the relative areas occupied by differentiating spermatogonia (i.e. type A_diff_ and B), but also by Leydig cells, was significantly larger than in the untreated control group; the area occupied by spermatozoa was still half that in controls indicating that the recovery did not yet reach this cell type (Figs. 1*A* and *B*).

**Fig. 1.**
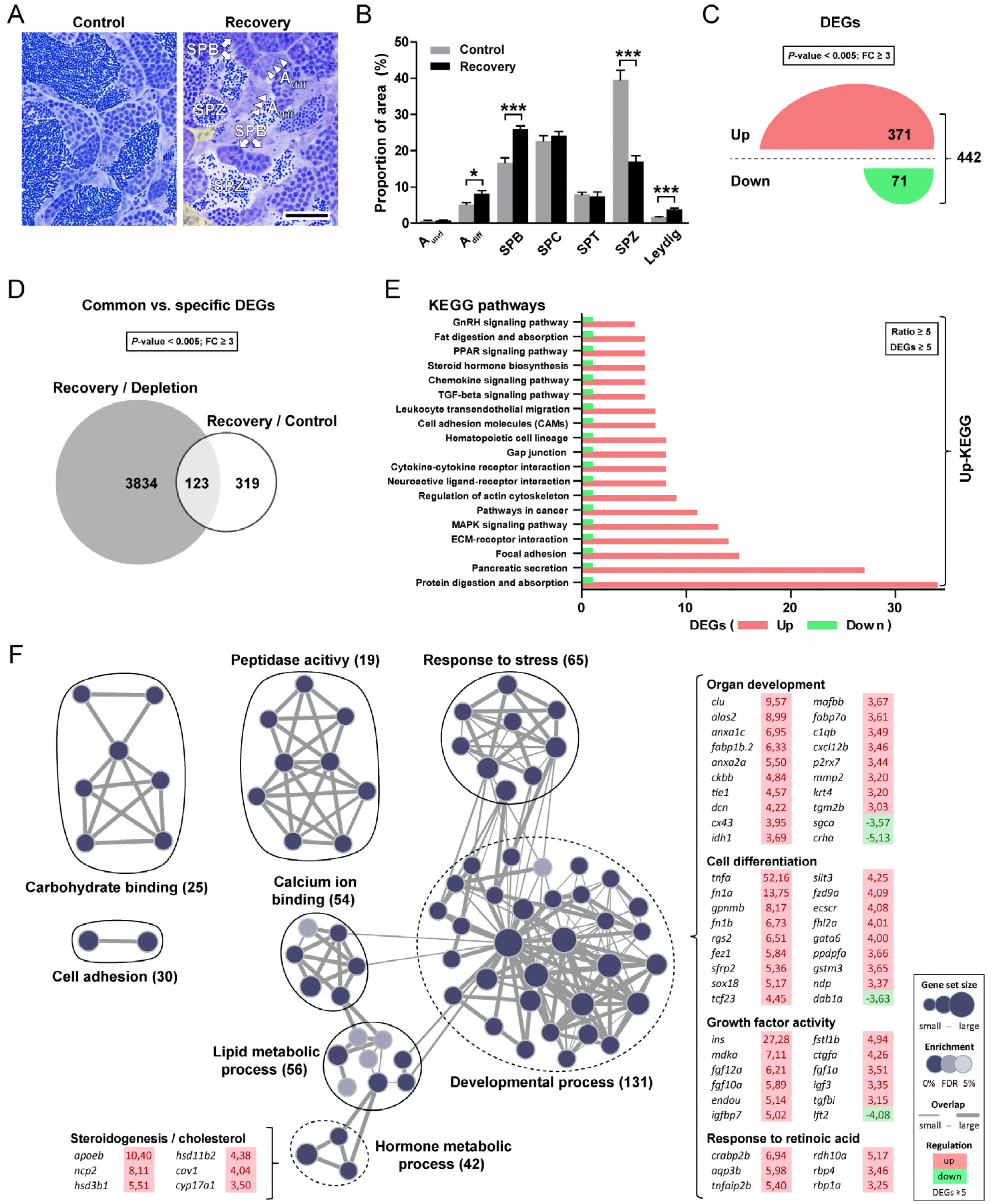
Morphometric and transcriptomic analyses of zebrafish testis during spermatogenic recovery from a cytotoxic (busulfan) insult. Qualitative (*A*) and quantitative (*B*) evaluation of the frequency of different germ and somatic cell types in recovering compared to untreated control testes. A_und_, type A undifferentiated spermatogonia; A_diff_, type A differentiating spermatogonia (arrowheads); SPB, type B spermatogonia (arrows); SPC, spermatocytes; SPT, spermatids; SPZ, spermatozoa (dashed line); Leydig, Leydig cells (yellow background, including other interstitial cells). Data are expressed as mean ± SEM (n = 5; * *P* < 0.05, *** *P* < 0.001, unpaired Student’s t test). Scale bar represents 25 μm. (*C*) Total numbers of up (red)- and down (green)-regulated genes identified by RNAseq at the beginning of the recovery period compared to the untreated control group (n = 5, *P* < 0.005, fold change [FC] ≥ 3.0, unpaired Student’s t test; see Dataset 1 for detailed information). (*D*) Common and specific recovery-associated genes retrieved when tested against the depleted (grey set) and control (white set) testis transcriptomes (n = 5, *P* < 0.005, [FC] ≥ 3.0, unpaired Student’s t test; see Dataset 1 for detailed information). (*E*) KEGG pathways modulated in recovering testes. Each pathway shown is represented by at least 5 DEGs and has a ratio of regulated genes (up-/down-, or *vice versa*) higher than 5. DEGs are highlighted with red (up-) or green (down-regulated) background. (*F*) Functional enrichment of recovery-induced gene expression in the zebrafish testis. RNAseq results were mapped (after Gene Ontology [GO] enrichment analysis) resulting in a network of functionally related gene sets (blue nodes) that form enrichment groups. Nodes represent statistically significant GO terms (*P* < 0.005, FDR < 0.05), links (grey lines) represent the number of overlapping genes (indicated by their thickness) between connected sets. Groups of closely related GO terms are encircled and labeled (numbers of regulated genes are shown). The groups labeled as Developmental process and Hormone metabolic process are highlighted with dashed lines and examples of identified DEGs in those sets are shown. **Figure 1-source data 1.** Raw data used to generate the statistical graphs in Figure 1.

In order to learn more about the molecular network controlling germ cell development during the recovery process, we carried out RNA sequencing (RNAseq) on: 1) untreated adult zebrafish testes, 2) germ cell-depleted testes, and 3) testis tissue collected at the beginning of the recovery period. Assuming that the recovery of spermatogenesis in our experimental model is achieved by stimulating both spermatogonial self-renewal and differentiation, we first checked for the expression of germ cell-marker genes. As expected, in depleted testes, in which A_und_ spermatogonia are the only germ cells present, we found strong expression of marker genes forSSCs and self-renewal factors (55), while the relative expression of these genes decreased in recovering testes that contain other germ cell types as well (*Supplementary file 1C*). In contrast, markers for differentiating spermatogonia were reduced in germ cell-depleted and elevated in recovering testis tissue (*Supplementary file 1C*). Comparison of the mRNA profiles of untreated and germ cell-depleted testes revealed that 2/3^rd^ of the differentially expressed genes (DEGs) were down- rather than up-regulated in depleted testes (1940 and 994, respectively; *Supplementary file 2A*). Similarly, the majority of KEGG terms significantly enriched in germ cell-depleted testes were down-regulated (*Supplementary file 2A*), including pathways related to cell cycle and meiosis, as well as others involved in a variety of development processes (i.e. Jak-STAT, Hedgehog, ErbB, or mTOR signaling). Among the up-regulated KEGG terms, steroid hormone biosynthesis, retinol metabolism and immune-related pathways were identified. Interestingly, most of the significantly modulated signaling pathways found in germ cell-depleted testes were also observed at the beginning of the recovery period but followed the opposite expression pattern (*Supplementary file 2A and B*). During recovery, the number of up-regulated DEGs exceeded the down-regulated ones (2344 and 1613, respectively; *Supplementary file 2B*), in contrast to the observation in depleted testes.

Analysis of the mRNA profiles of recovering and control testes provided a much lower number of DEGs (i.e. 442; Fig. 1*C*) than in the previous comparison (i.e. 3957; *Supplementary file 2B*), but showed a similar pattern since most transcripts identified were stimulated in their expression levels (371 out of 442; Fig. 1*C*). All KEGG pathways significantly affected were up-regulated (Fig. 1*E*), including TGF-beta, steroid hormone, GnRH, PPAR, MAPK signaling systems, as well as extracellular matrix (ECM) structure and remodeling-related pathways (such as focal adhesion, cell adhesion molecules, ECM-receptor interaction). When searching for common and specific DEGs, 123 genes were found in both data sets and 3834 or 319 specifically regulated during recovery with respect to the depleted or control conditions, respectively (Fig. 1*D*). Examining the biological functions enriched among recovery-regulated genes by GO analysis showed numerous overlapping gene sets (sharing a similar function/GO identifier) that were grouped in 8 different clusters (encircled in Fig. 1*F*). 131 DEGs form the main cluster (Developmental process; Fig. 1*F*), which includes genes associated with organ development (e.g. *clu*, *dcn*, *cx43*, *mmp2*, *krt4*, *cxcl12b*, *fabp7a*), cell differentiation (e.g. *tnfa*, *tcf23*, *ecscr*, *sox18*, *ppdpfa*, Wnt-related [*sfrp2*, *fzd9a*, *ndp*]), growth factor activity (e.g. insulin-like [*ins*, *igfbp7*, *igf3*], TGF-beta [*fstl1b*, *tgfbi*, *lft2*], and FGF [*fgf1a*, *fgf10a*, *fgf12a*] family members) and retinoic acid signaling (e.g. *crabp2b*, *rdh10a*, *rbp1a*, *rbp4*). Furthermore, gene sets related to a variety of different metabolic processes were enriched in recovering compared to untreated control testes (Lipid metabolic process, Response to stress, Calcium ion binding and Hormone metabolic process clusters; Fig. 1*F*). Within the list of DEGs belonging to that category, steroidogenesis-related enzymes were increased in their transcript levels (e.g. *hsd3b1*, *hsd11b2* and *cyp17a1*). Not connected to the main group, or to each other, functional enrichment results moreover include clusters such as Cell adhesion, Carbohydrate binding and Peptidase activity (Fig. 1*F*).

### Retinoid-mediated stimulation of spermatogonial development

Having found several genes involved in RA-signaling, we studied the potential role of this signaling system in zebrafish spermatogenesis. First, we examined if the significant up-regulation of the GO-term “Response to retinoic acid” could be supported experimentally, using a primary testis tissue culture system. Upon *ex vivo* exposure to the RA precursor RE, the area in testis sections occupied by type A_diff_ and B spermatogonia increased (Figs. 2*A* and *C*), as well as the proliferation activity of these germ cell types (Figs. 2*B* and *D*). Consistent with the morphological data, the mRNA levels of *piwil1* and *dazl* increased (Fig. 2*E*). Also, elevated transcript levels of the RA-producing enzyme *aldh1a2* and of *fshr* were observed, but exposure to RE did not modulate the transcript levels of selected growth factors (*amh*, *igf3*, *insl3*; Fig. 2*E*). Similar changes in germ cell-marker gene expression were found in response to RE and RA when examining primary cultures of enzymatically dispersed testicular cells, with elevated *piwil1* and *nanos2* transcript levels (*Supplementary file 3A*). Also, *aldh1a2* expression was stimulated by RE in both cell and tissue cultures (Fig. 2*E* and *Supplementary file 3A*), while RA did not have this effect in cell suspension experiments (*Supplementary file 3A*). In contrast, while RE did not change *cyp26a1* mRNA (encoding an enzyme metabolizing RA) levels in tissue cultures, it did so in cell cultures, an effect that was even stronger after exposure to RA (*Supplementary file 3A*). Consistent with the RE-induced increase in type A_diff_ and B spermatogonia (Fig. 2*C*), chemical inhibition of RA production by DEAB decreased the frequency of these spermatogonia in tissue culture (Fig. 2*F*). The DEAB-induced increase in germ cell apoptosis may have contributed to the decrease in the section area occupied by differentiating spermatogonia (Fig. 2*F*), as DEAB treatment did not affect spermatogonial proliferation (Fig. 2*G*). In terms of gene expression, slight but significant decreases in *fshr* transcripts levels were found in the presence of DEAB, while growth factor expression did not change (Fig. 2*H*). These data indicate that RA signaling promotes differentiating spermatogonial numbers by stimulating their proliferation and reducing germ cell apoptosis

**Fig. 2.**
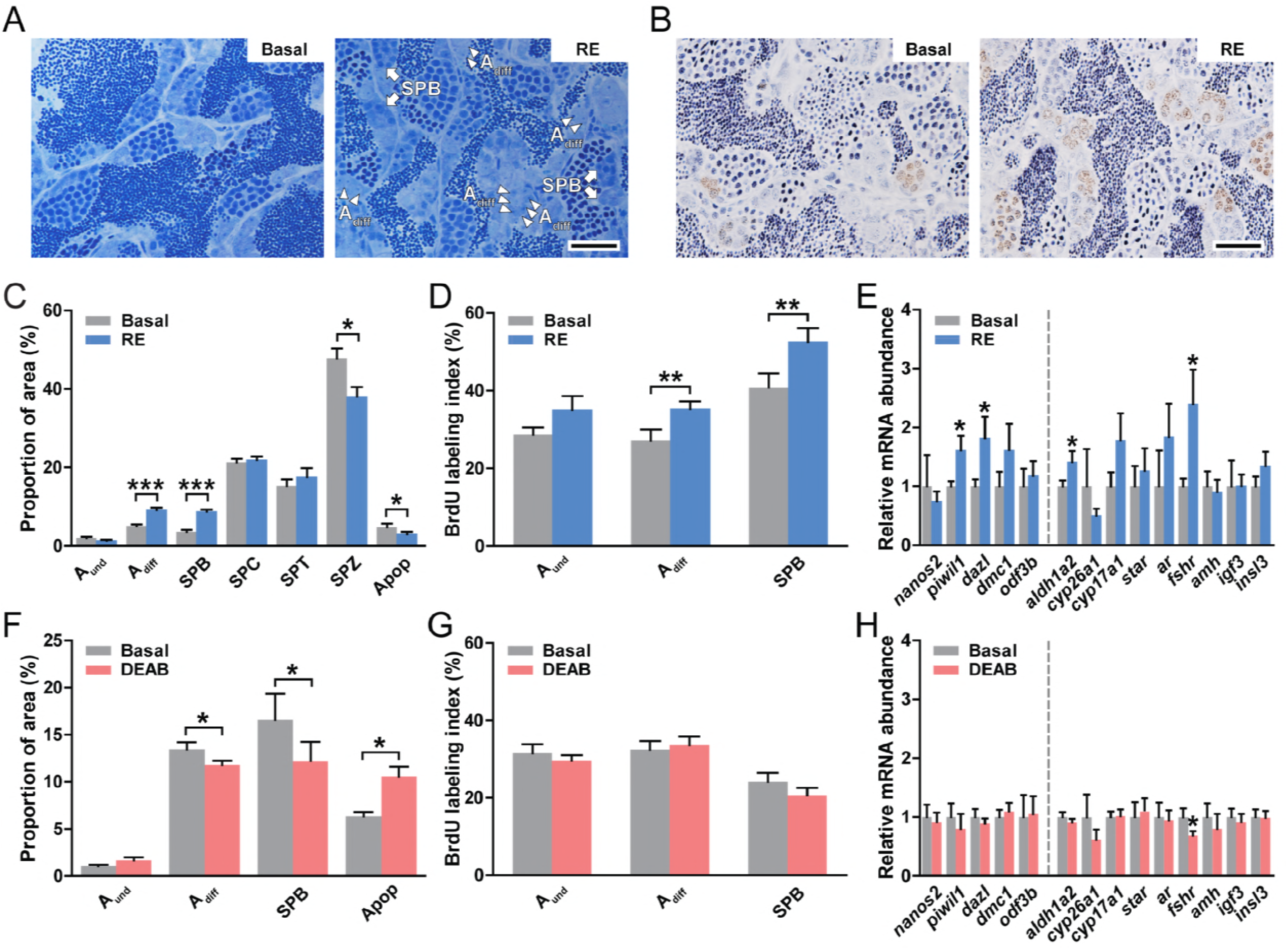
Retinoid effects on spermatogonial development. Qualitative (*A* and *B*) and quantitative evaluation of the proportions of different germ cells and apoptotic cells (*C* and *F*), of the proliferation activity of spermatogonia (*D* and *G*), and of the transcript levels of selected genes (*E* and *H*) in testes cultured for 4 days under different experimental conditions: in the absence or presence of retinol (RE, 10 μM; *C* and *D*), and in the absence or presence of DEAB (10 μM; *F* and *G*). A_und_, type A undifferentiated spermatogonia; A_diff_, type A differentiating spermatogonia (arrowheads in *A*); SPB, type B spermatogonia (arrows in *A*); SPC, spermatocytes; SPT, spermatids; SPZ, spermatozoa; Apop, apoptotic cells. Data are expressed as mean ± SEM (n = 6; * *P* < 0.05, ** *P* < 0.01, *** *P* < 0.001, paired Student’s t test). Scale bar represents 25 μm. In *E* and *H*, data are shown as mean of fold change ± SEM (n = 7-12) and expressed relative to the control condition, which is set at 1. Asterisks indicate significant differences between treated and control groups (* *P* < 0.05. paired Student’s t test). **Figure 2-source data 1.** Raw data used to generate the statistical graphs in Figure 2.

### *sall4* and *rec8* mediate RA effects in the zebrafish testis

To find the mechanisms mediating RA effects in the zebrafish (missing a *stra8* gene), we tested the expression of *sall4* and *rec8* (*a* and *b* paralogs), previously identified as components of *Stra8*-independent pathways mediating RA signaling in mammals (5, 6). We found that RA and RE increased the transcript levels of *sall4* and *rec8* (Fig. 3*A*). We then examined if knocking down *sall4*, *rec8a* and *rec8b* transcripts mimicked the effects of a DEAB-induced reduction of RA levels on germ cell development. Specific GapmeRs for *sall4*, *rec8a* and *rec8b* reduced target transcripts levels and reduced the expression of *piwi1l* and *dazl*, marker genes for differentiating spermatogonia, and also for meiotic markers *sycp3* and *odf3b* (Figs. 3 *B*-*D*). In order to investigate the cellular expression of *sall4*, *rec8a* and *rec8b*, pooled testicular cell suspensions obtained from *Tg(vasa:EGFP)* testes (Fig. 3*E*) were FACS sorted and EGFP-negative and EGFP-positive fractions were analyzed by qPCR. As shown in Fig. 3*F*, *sall4* transcripts were enriched in EGFP-positive cells, indicating its preferential expression in germ cells (see *Supplementary file 3B* for further details). In addition, *sall4* expression was reduced in busulfan-treated testes (−6.59-fold; *Dataset 1*). In a similar way, *rec8* paralogues tended to be enriched in the EGFP-positive fraction (Fig. 3*F*). Altogether, these observations suggest that at least in part retinoids stimulate zebrafish spermatogenesis in a *sall4*- and *rec8*-dependent manner through factors operating in germ cells.

**Fig. 3.**
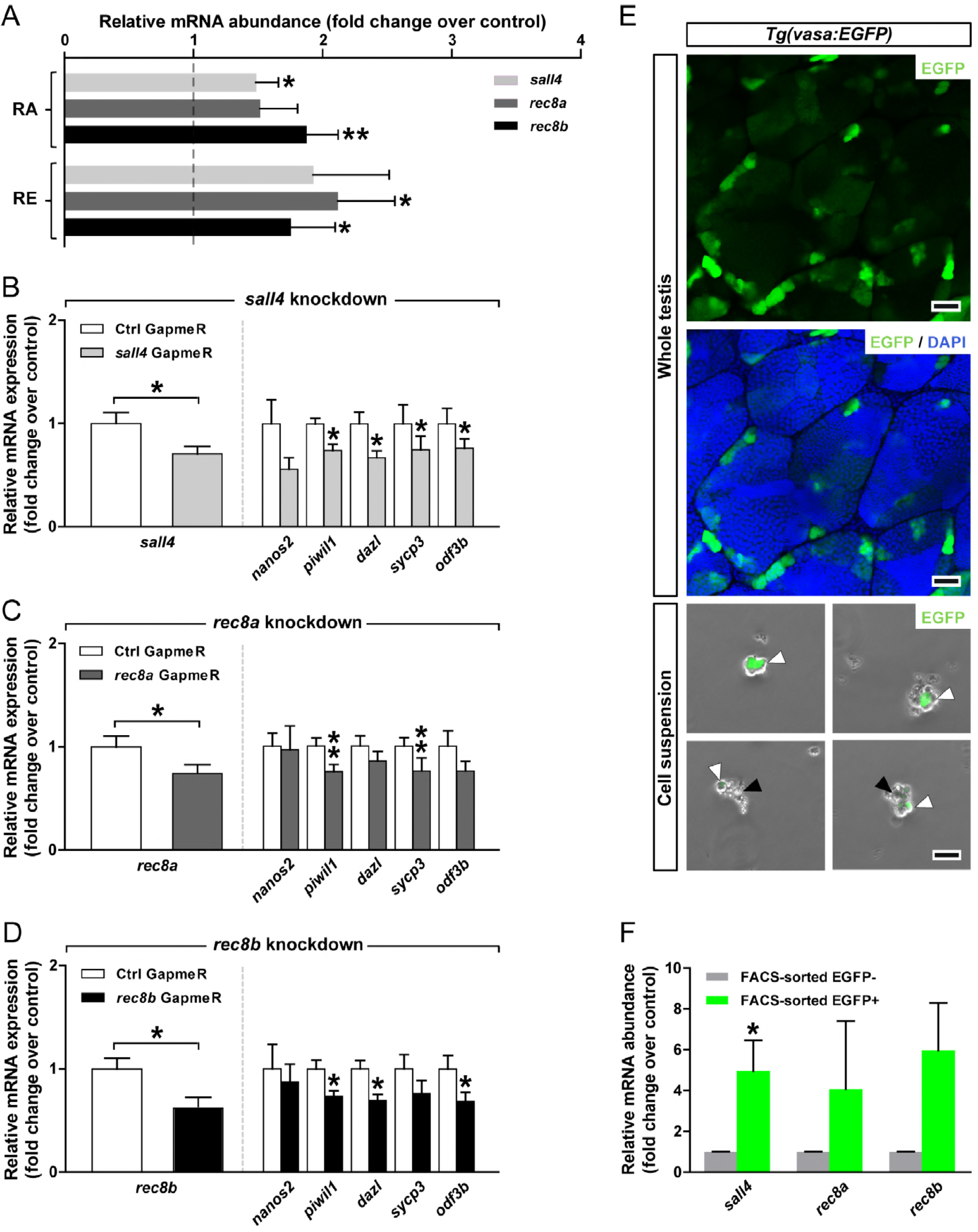
Involvement of *sall4*, *rec8a* and *rec8b* in mediating RA effects in the zebrafish testis. (*A*) Modulation of *sall4*, *rec8a* and *rec8b* mRNA levels in response to retinoids. Testicular explants were cultured for 4 days with RA (10 μM) or RE (10 μM). Data are expressed as mean fold change ± SEM (n = 6-15; * *P* < 0.05, ** *P* < 0.01, paired Student’s t test) and shown as relative to the control condition, which is set at 1 (dashed line). (*B*-*D*) Effects of GapmeR (1 μM)-mediated knockdown of *sall4*, *rec8* or *rec8b* after 4 days of tissue culture, on the targeted and selected germ cell-marker gene transcripts. Data are shown as mean fold change ± SEM (n = 8-13, including three technical replicates; * *P* < 0.05, ** *P* < 0.01, paired Student’s t test) and normalized to the GapmeR control (Ctrl), which is set at 1. (*E*) A pooled testicular cell suspension was obtained from 20 *Tg(vasa:EGFP)* testes and subjected to FACS (see *Supplementary file 3B* for further details). Black and white arrowheads indicate EGFP^-^ and EGFP^+^ cells, respectively. (*F*) Relative mRNA expression of *sall4*, *rec8a* and *rec8b* in FACS-sorted EGFP^-^ and EGFP^+^ fractions. Data are shown as mean fold change ± SEM (n = 3, technical replicates; * *P* < 0.05, unpaired Student’s t test) and expressed relative to the EGFP^-^ condition, which is set at 1. **Figure 3-source data 1.** Raw data used to generate the statistical graphs in Figure 3.

### A dominant-negative *raraa* transgene targeted to germ cells disturbs spermatogenesis

To obtain further evidence for the germ cell-specific role(s) of RA-signaling, we generated transgenic zebrafish (named *dn-raraa*) expressing a truncated form of the RA receptor alpha a (Raraa) in germ cells, in a similar way as described previously for mammals (56) and zebrafish (57).

Transgenesis efficiency was first analyzed in larvae at 3-4 days post-fertilization, based on EGFP-positive hearts and mCherry-positive germ cells in the genital ridge (Fig. 4*A*). Transgene expression was confirmed in a later stage as shown in *Supplementary file 4A*. Morphological evaluation of 6 months adult *dn-raraa* testes revealed clear defects in spermatogenesis, including germ cell-depleted areas, abnormal cystic organization and massive apoptosis (Fig. 4*Biii-vi* and *Supplementary file 4B*). Quantitative evaluation of spermatogenesis showed relative smaller areas occupied by type A_diff_ spermatogonia and spermatocytes. The category “Others” (including empty areas and germ cell-depleted tissue), was also higher in 6 months adult *dn-raraa* testes (Fig. 4*C*). Unexpectedly, the germinal epithelium had largely recovered in 9 months adult *dn-raraa* males (Figs. 4*B*vii-viii and *C*), even showing a higher gonadosomatic index than wild type (WT) siblings (Fig. 4*D*). Despite the apparent recovery of spermatogenesis, clear fertilization problems were associated with the over-expression of the truncated receptor encoded by the *raraa^DN391^* transgene, since the ability to fertilize eggs was low in all transgenic compared to WT males, which may be related to the increased apoptotic incidence of post-meiotic germ cells found in 9 months-old transgenic males (Fig. 4*Bix-xi*). Moreover, those eggs that were fertilized with transgenic sperm showed a four times lower survival rate (Fig. 4*E*). Analysis of RA-related gene expression showed a consistent down-regulation of *rec8a* in adult *dn-raraa* males, while the *raraa^DN391^* transgene mRNA levels were strongly enhanced compared to WT (undetectable levels in all WT testis samples; Fig. 4*F*). Transcript levels of steroid production/signaling genes (*star*, *hsd3b1* and *ar*) were significantly enhanced, but only in the 6 months-old *dn-raraa* males.

**Fig. 4.**
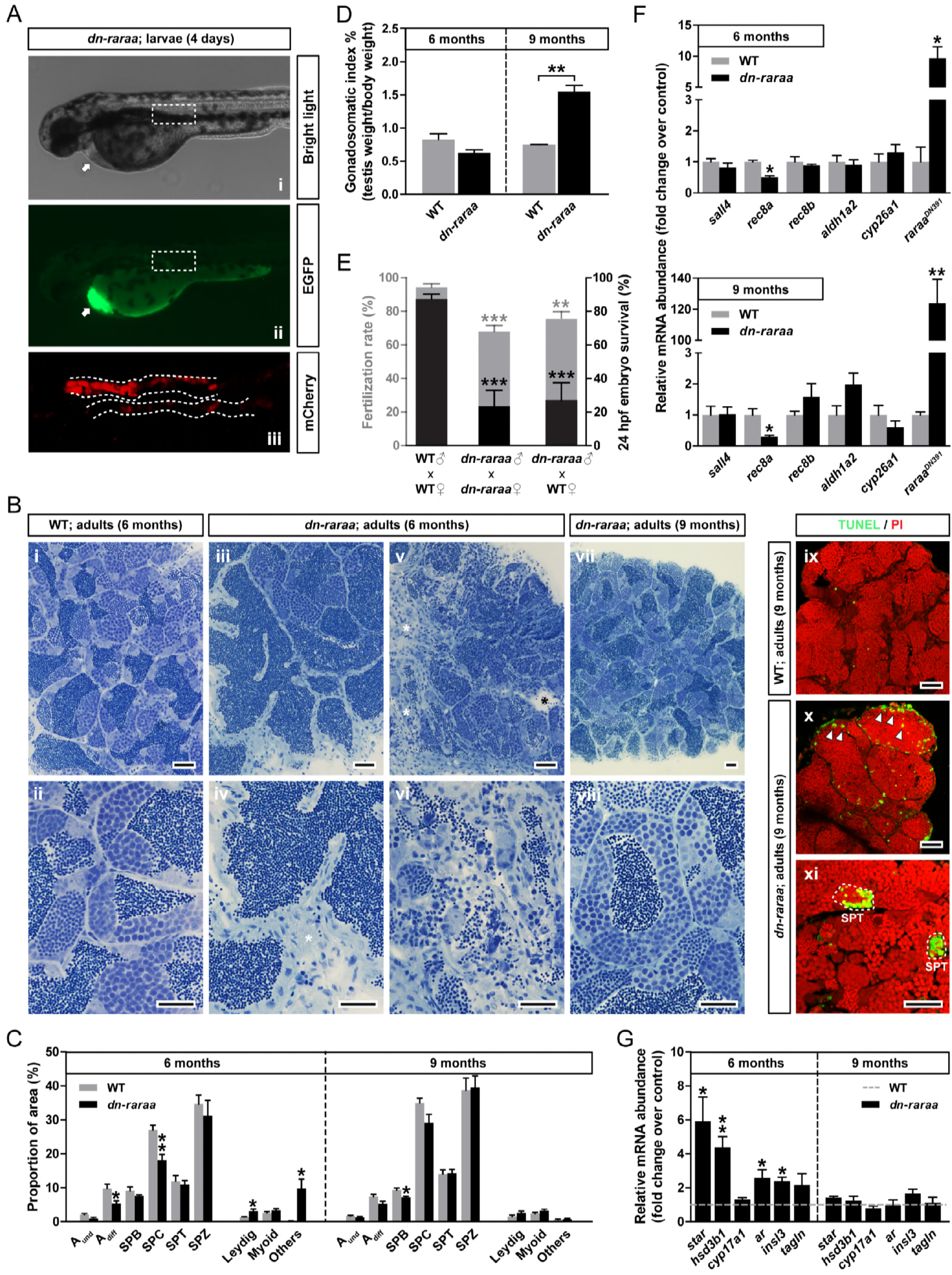
Inactivation of retinoid signaling in germ cells results in severe testicular defects. (*A*) Confirmation of transgenesis 4 days post-fertilization in zebrafish larvae microinjected as 1 cell stage embryos. EGFP and mCherry expression was detected in the heart and genital ridge of *dn-raraa* larvae, respectively. (*B*-*D*) Qualitative (*B*) and quantitative analysis of spermatogenesis (*C*) and gonadosomatic indices (*D*) of WT and *dn-raraa* males after 6 and 9 months post-fertilization (data expressed as mean ± SEM; n = 3-5; * *P* < 0.05, ** *P* < 0.01, unpaired Student’s t test). Scale bar represents 25 μm. White asterisks denote germ cell-depleted tissue while black asterisks indicate empty areas. A_und_, type A undifferentiated spermatogonia; A_diff_, type A differentiating spermatogonia; SPB, type B spermatogonia; SPC, spermatocytes; SPT, spermatids; SPZ, spermatozoa; Leydig, Leydig cells; Myoid, myoid cells; Others, germ cell-depleted tissue plus empty areas. In *B* (panels *ix*-*xi*), confirmation of germ cell apoptosis by TUNEL analysis is presented. TUNEL-positive cells/cysts are shown in green and PI (propidium iodide) counterstain in red. Arrowheads indicate isolated apoptotic cells among spermatozoa in the tubular lumen and representative spermatid cysts containing several apoptotic cells are encircled with a dashed line. Scale bar represents 25 μm. (*E*) Fertilization rate (grey bars) and embryo survival of fertilized eggs (black bars) from adult WT and *dn-raraa* transgenic males. Mating was repeated every 7 days (data are shown as mean ± SEM; n = 4-6; ** *P* < 0.01, *** *P* < 0.001, one-way ANOVA followed by Tukey’s multiple comparison test) and fertilization and survival recorded 2 and 24 hours post-fertilization (24 hpf), respectively. (*F*) qPCR quantification of RA-related gene and *raraa^DN391^* transgene transcripts in 6 and 9 months-old WT and *dn-raraa* testicular samples. Data are shown as mean fold change ± SEM (n = 3-4; * *P* < 0.05, ** *P* < 0.01, unpaired Student’s t test) and expressed relative to the WT group, which is set at 1. (*G*) Transcript levels of steroidogenesis-related (*star*, *hsd3b1*, *cyp17a1* and *ar*), Leydig cell (*insl3*) and myoid cell (*tagln*) genes in WT and *dn-raraa* transgenic adult testes. Data are shown as mean fold change ± SEM (n = 3-4; * *P* < 0.05, ** *P* < 0.01, unpaired Student’s t test) and expressed relative to the WT group, which is set at 1 (dashed line). **Figure 4-source data 1.** Raw data used to generate the statistical graphs in Figure 4.

Interestingly, the relative area occupied by Leydig cells and the expression levels of the Leydig cell-derived factor *insl3* increased (but not the myoid cell-specific *tagln*; Fig. 4*G*). Therefore, our observations suggest that Leydig cell products are involved in promoting the spermatogenic recovery in the absence of germ cell RA signaling. This recovery process had reached a new equilibrium at 9 months of age, allowing full spermatogenesis although sperm quality remained poor.

### RA supports androgen production

Since disturbing RA signaling in germ cells promoted Leydig cell activity, which usually is regulated by pituitary gonadotropins, we speculated that in our *dn-raraa* mutant males, a feedback loop involving the brain-pituitary system became activated. We therefore decided to examine RA effects on basal and gonadotropin-stimulated testicular androgen production. As gonadotropin, we used zebrafish Fsh, which is a potent steroidogenic hormone as not only Sertoli but also Leydig cells express the Fsh receptor; moreover, only Fsh, but not Lh, increased the expression of selected steroidogenic genes (47).

Fsh has a clear stimulatory effect on androgen production. However, when RA synthesis was inhibited after 2 days of tissue culture, the Fsh-induced stimulation on androgen release was compromised, and could not be reversed by adding exogenous RA (Fig. 5*A*). While basal androgen release was not changed after 2 days (Fig. 5*A*), it did decrease after inhibiting RA production when the incubation period was prolonged to 7 days (Fig. 5*B*). Further evidence for the relevance of RA signaling for steroid production was obtained by GapmeR-mediated knockdown of *sall4* and *rec8a* that reduced testicular transcript levels of the steroidogenesis-related genes *star* and *cyp17a1* (Figs. 5*C* and *D*). Knockdown of *rec8b* had no effect (data not shown).

**Fig. 5.**
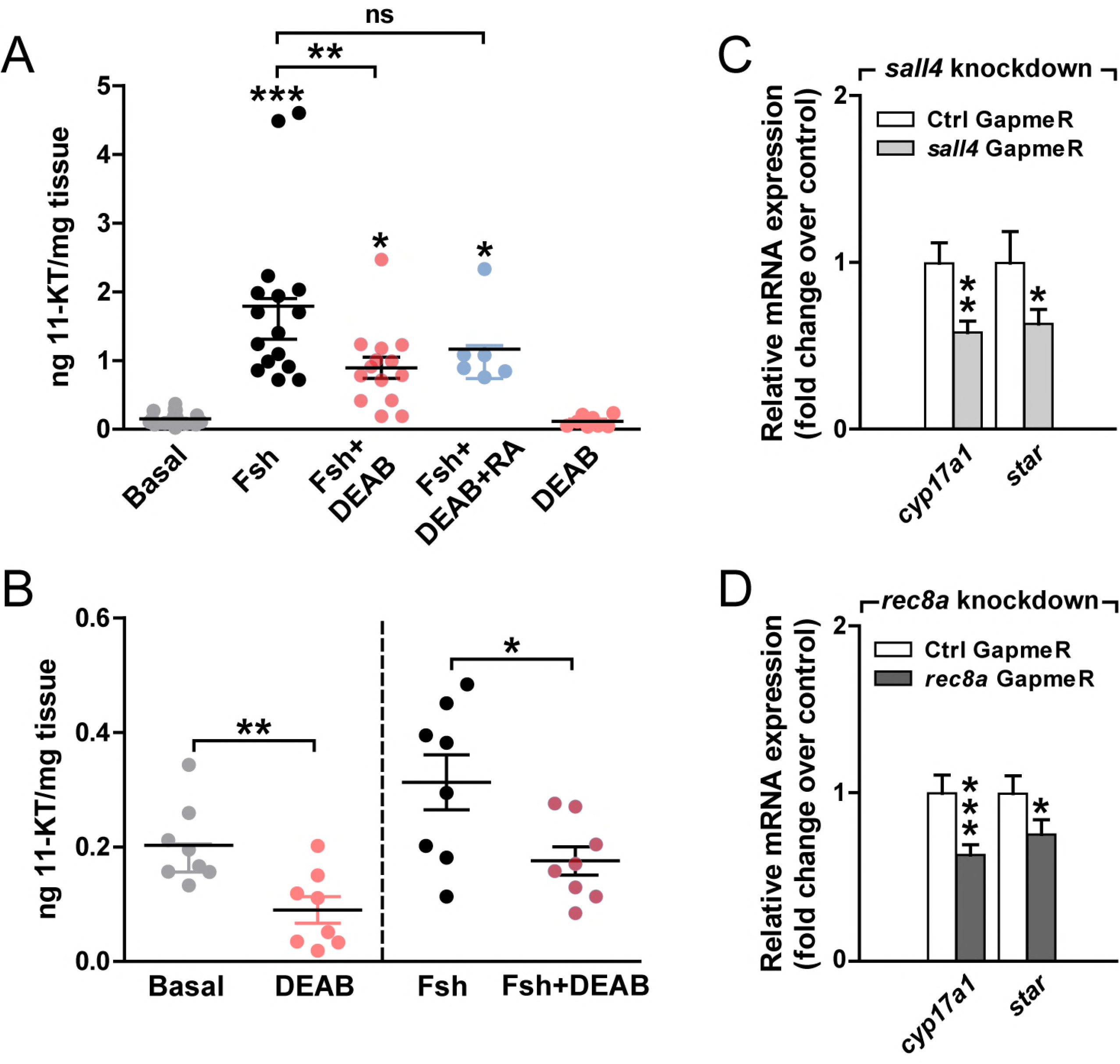
Inhibition of retinoid signaling alters testicular androgen production and steroidogenesis-related gene expression. (*A* and *B*) Quantification of 11-ketotestosterone (11-KT) production by testis tissue cultured for 2 (*A*) or 7 (*B*) days in response to 100 ng/mL Fsh, 10 μM DEAB, or 10 μM RA. 11-KT levels were determined by RIA. Data are expressed as mean ± SEM (n = 6-20; * *P* < 0.05, ** *P* < 0.01, *** *P* < 0.001, one-way ANOVA followed by Tukey’s multiple comparison test (*A*) or paired Student’s t test (*B*)). “ns” denotes no significant differences. (*C* and *D*) qPCR results of effects of *sall4* (*C*) and *rec8a* (*D*) knockdown via specific GapmeRs (1 μM) on steroidogenesis-related gene expression in tissue culture conditions for 4 days. Data are shown as mean fold change ± SEM (n = 8-12, including three technical replicates; * *P* < 0.05, ** *P* < 0.01, *** *P* < 0.001, paired Student’s t test) and normalized to the GapmeR control (Ctrl), which is set at 1. **Figure 5-source data 1.** Raw data used to generate the statistical graphs in Figure 5.

### Androgen and RA signaling interact to drive spermatogenesis

Our data suggest that RA signaling supports androgen production, as was reported very recently in mice (58). However, it is not known whether RA also supports stimulation of spermatogenesis by androgens. To study this, we first examined androgen effects alone. Testis tissue responded to 11-ketotestosterone (11-KT), the main teleost androgen, by an up-regulation of several germ cell-marker genes (Figs. 6*A* and *B*). Morphometric evaluation showed that 11-KT slightly increased the numbers of A_diff_ spermatogonia and more clearly of spermatids, while reducing the incidence of apoptosis (Fig. 6*C*). Inhibiting RA production reduced the stimulatory effect of 11-KT on 4 out of 7 germ cell-markers (Figs. 6*D* and *E*), which was also reflected in reduced proportions of spermatocytes and spermatids, the latter just not reaching significance (Fig. 6*F*). Surprisingly, the presence of 11-KT could not prevent the DEAB-induced increase in apoptosis (Fig. 6*F*). TUNEL analysis confirmed that inhibiting RA synthesis increased the incidence of DNA fragmentation, mainly in spermatids but also in spermatocytes (Figs. 6*G*, *H* and *Supplementary file 5*, identified by the shape and size of their DAPI-stained nuclei), despite the presence of androgens. Considering that androgens decreased germ cell apoptosis (Fig. 6*C*) and that blocking RA production increased apoptosis irrespective of the absence (Fig. 2*F*) or presence (Fig. 6*F*) of androgens, we wondered if inhibiting steroid synthesis in the presence of RE affected apoptosis. Since this was not the case (Fig. 7*A*), it seems possible that androgens elevate RA production, which may then prevent/reduce apoptosis. Indeed, short-term (4 days) *ex vivo* and long-term (5 weeks) *in vivo* exposure to androgen increased *aldh1a2* transcript levels; also *cyp26a1* transcript levels increased, although not significantly in the *in vivo* experiment (Fig. 7*D*). These results suggest that androgen-induced RA production and metabolism reduces apoptotic germ cell loss.

**Fig. 6.**
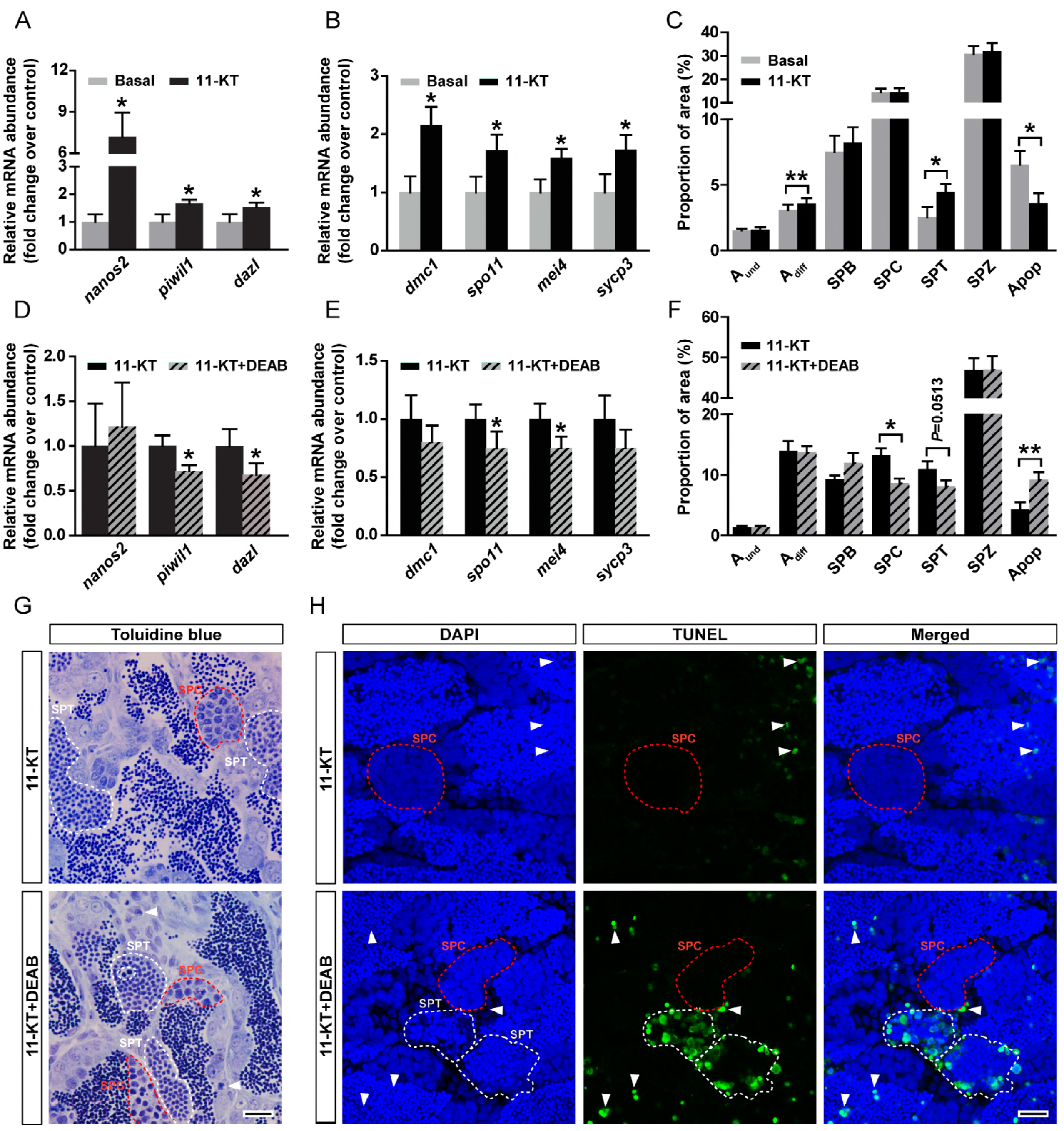
Retinoid involvement in androgen-stimulated spermatogenesis. (*A*-*C*) Transcript levels of selected spermatogonial (*A*) and meiotic (*B*) markers, and frequency of different germ cell types and apoptotic cells (*C*) in testes incubated for 4 days in basal conditions or with 200 nM 11-KT. Data are shown as mean of fold change ± SEM (n = 8; *, *P* < 0.05, ** *P* < 0.01, paired Student’s t test) and expressed relative to the respective control condition, which is set at 1. (*D*-*H*) Effects of inhibiting RA production on androgen-stimulated spermatogenesis. Transcript levels of germ cell-marker genes (*D* and *E*) and frequency of different germ cell types and apoptotic cells (*F*) in testes incubated for 4 days with 200 nM 11-KT, in the absence or presence of DEAB (10 μM). Data are expressed as mean ± SEM (n = 7-8; * *P* < 0.05, ** *P* < 0.01, paired Student’s t test). (*G*) Germ cell apoptosis in testicular explants incubated with 11-KT in the absence or presence of DEAB. Arrowheads indicate isolated apoptotic cells and representative spermatocyte/spermatid (apoptotic or not) cysts are encircled with a red or white dashed line, respectively. (*H*) Confirmation of germ cell apoptosis by TUNEL analysis. TUNEL-positive cells/cysts are shown in green and DAPI counterstain in blue. Arrowheads indicate isolated apoptotic cells and representative spermatocyte/spermatid (TUNEL^+^ or TUNEL^-^) cysts are encircled with a red or white dashed line, respectively. A_und_, type A undifferentiated spermatogonia; A_diff_, type A differentiating spermatogonia; SPB, type B spermatogonia; SPC, spermatocytes; SPT, spermatids; SPZ, spermatozoa; Apop, apoptosis. Scale bar represents 15 μm in *G* and 10 μm in *H*. **Figure 6-source data 1.** Raw data used to generate the statistical graphs in Figure 6.

**Fig. 7.**
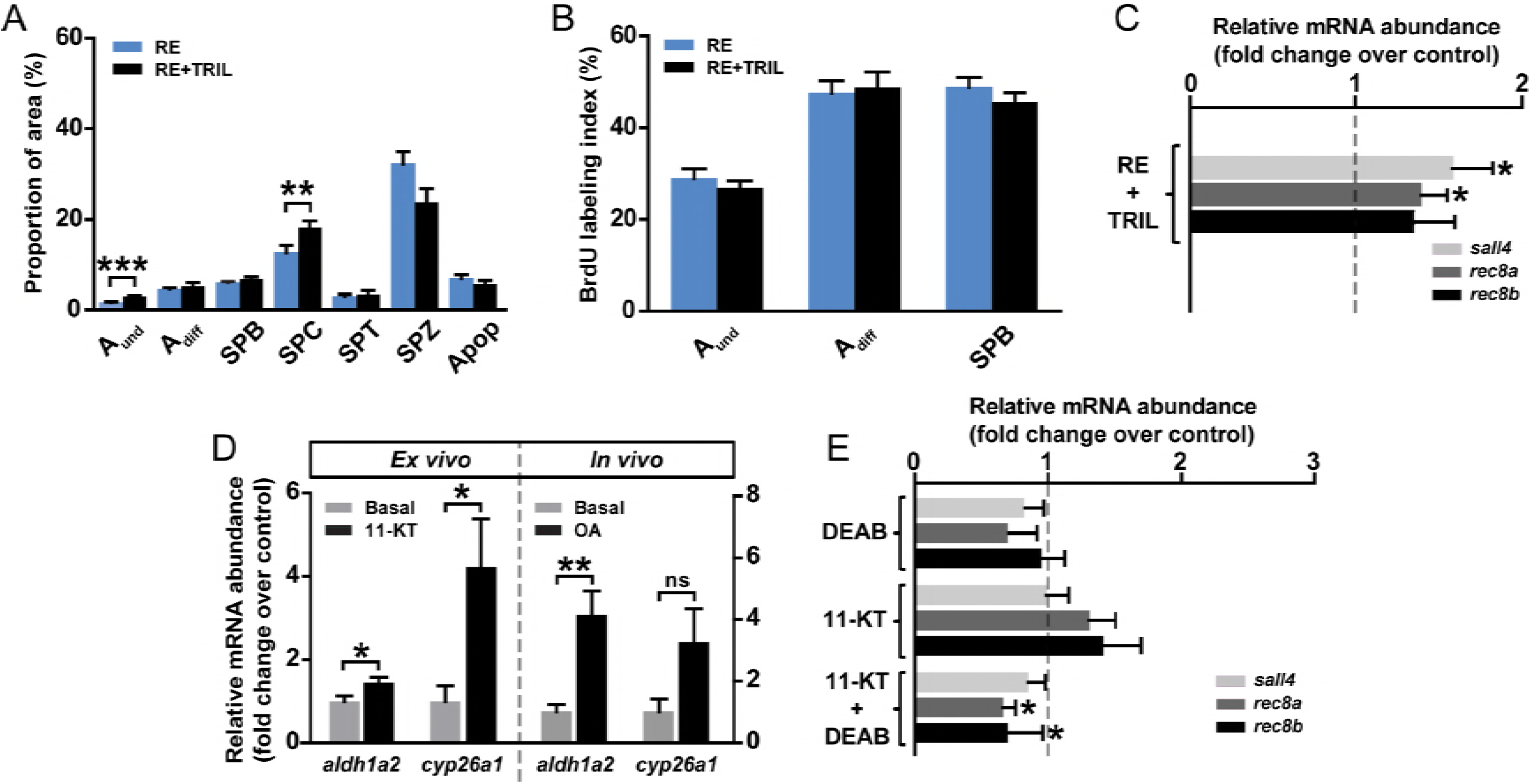
Steroid influence on the regulation of retinoid-modulated spermatogenesis. Frequency of germ cell types and apoptotic cells (*A*), BrdU labeling indices (*B*) and RA target genes (*C*) in testicular explants cultured for 4 days in the presence of 10 μM RE or in addition of 25 μg/mL trilostane (TRIL). Data are expressed as mean ± SEM (n = 6-12; * *P* < 0.05, ** *P* < 0.01, *** *P* < 0.001, paired Student’s t test). A_und_, type A undifferentiated spermatogonia; A_diff_, type A differentiating spermatogonia; SPB, type B spermatogonia; SPC, spermatocytes; SPT, spermatids; SPZ, spermatozoa; Apop, apoptosis. In *C*, results are expressed relative to the basal control group, which is set at 1 (dashed line), and RE-induced levels are shown in Fig. 3*A*. (*D*) *Ex vivo* and *in vivo* androgen effects on *aldh1a2* and *cyp26a1* expression. In the left panel, testis tissue was cultured for 4 days to study 11-KT (200 nM) effects on the mRNA abundance of RA-related enzymes compared to the control group. In the right panel, adult zebrafish males were exposed to 100 nM 11-ketoandrostenedione (OA) *in vivo* for 5 weeks. Data are expressed as mean fold change ± SEM (n = 7-8; * *P* < 0.05, ** *P* < 0.01, paired (*ex vivo*) or unpaired Student’s t test (*in vivo*)) and shown as relative to the control condition, which is set at 1. “ns” denotes no significant differences. (*E*) RA target gene expression in testicular explants cultured for 4 days under basal or 11-KT (200 nM)-stimulated conditions, in the absence or presence of 10 μM DEAB. Data are expressed as mean fold change ± SEM (n = 7-8; * *P* < 0.05, paired Student’s t test) and shown as relative to the basal condition, which is set at 1 (dashed line). **Figure 7-source data 1.** Raw data used to generate the statistical graphs in Figure 7.

Inhibiting steroid production in the presence of RE also provided evidence for a so far undetected steroid effect: A_und_ spermatogonia accumulated in the absence of steroids (Fig. 7*A*) without affecting their proliferative activity (Fig. 7*B*). This suggests that in zebrafish steroid hormones stimulate the transition of A_und_ to A_diff_ spermatogonia. Considering that the RE-induced changes of *sall4* and *rec8a* transcript levels remained unaffected when inhibiting steroid production by trilostane (TRIL; Fig. 7*C*), it seems unlikely that steroids modulate retinoid effects but may instead support RA production (see Figs. 7 *D* and *E*). Similarly, the observed accumulation of spermatocytes (Fig. 7*A*) suggests that steroid hormones stimulate spermatocyte development. This possibility is in line with the androgen-induced increase in spermatids numbers (Fig. 6*C*).

### Retinoid signaling contributes to Fsh-stimulated spermatogonial differentiation

Basal and Fsh-stimulated androgen production as well as androgen action were supported by RA-signaling (Figs. 5 and 6), and it is possible that androgens stimulated RA production/metabolism (Fig. 7). Since Fsh is a major regulator of androgen production in fish, we asked if Fsh –via steroid hormones or otherwise– modulates testicular RA production.

Blocking RA production reduced the Fsh-stimulated proliferative activity of A and B spermatogonia, decreased the frequency of type A_diff_ and B (but not A_und_) spermatogonia, and increased the incidence of apoptosis among germ cells (Figs. 8*A* and *B*). Interestingly, Fsh increased testicular RA production (Fig. 8*C*) and subsequently the expression of the RA target genes *sall4*, *rec8a* and *rec8b*. This stimulation occurred in a steroid-independent manner but was blocked by inhibition of RA synthesis (Fig. 8*D*). This suggests that part of the biological activity of Fsh is mediated via the stimulation of RA production.

**Fig. 8.**
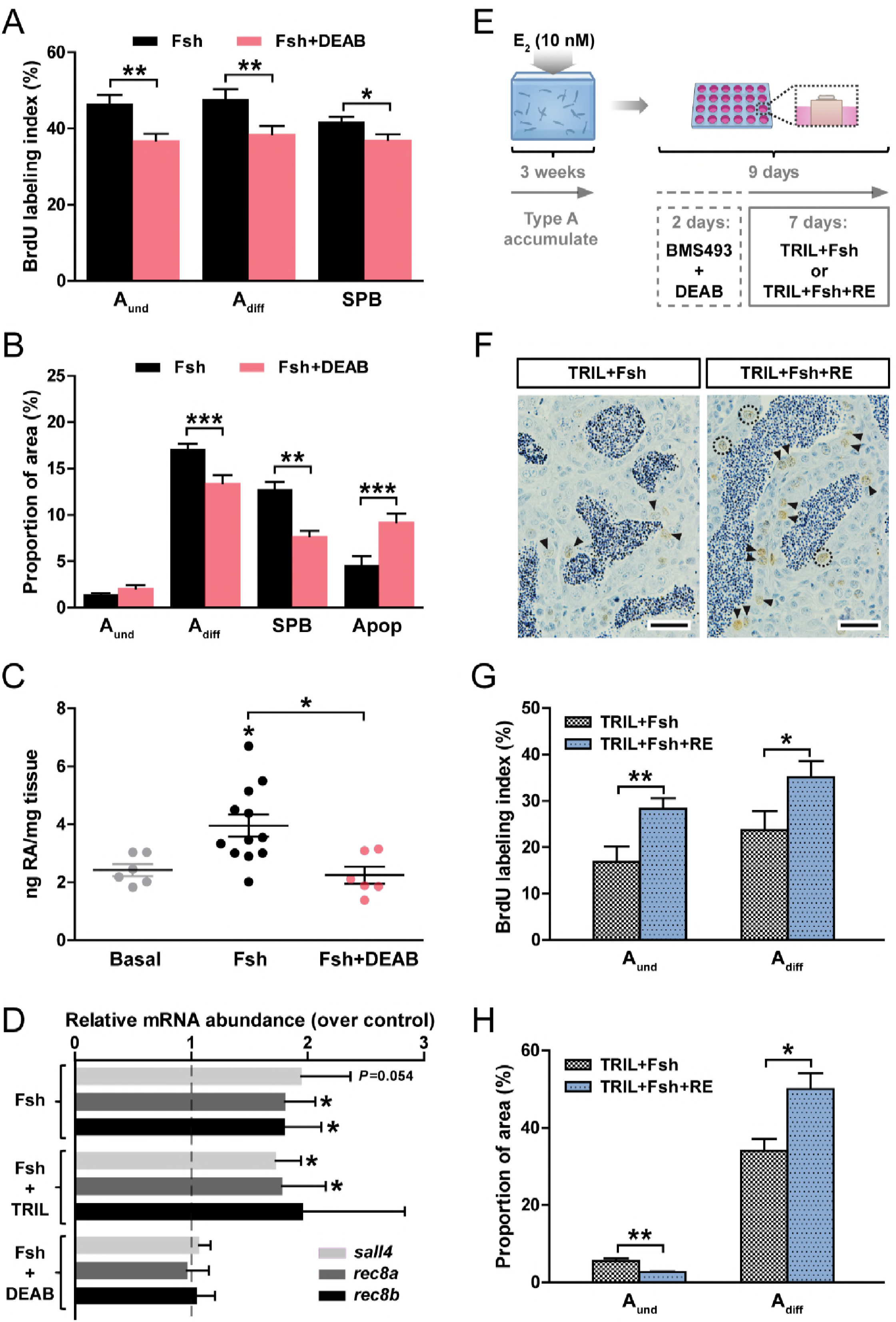
RA signaling contributes to Fsh-stimulated spermatogonial differentiation. Determination of BrdU labeling indices (*A*) and of frequency (*B*) of type A and B spermatogonia and apoptotic cells. Testicular explants were cultured for 4 days under Fsh-stimulated (100 ng/mL) conditions, in the absence or presence of 10 μM DEAB. (*C*) Quantification of retinoic acid (RA) production by testis tissue cultured for 4 days in response to basal medium, 100 ng/mL Fsh alone or with 10 μM DEAB. (D) Expression of RA target genes in testicular explants cultured for 4 days under Fsh-stimulated (100 ng/mL) conditions, in the absence or presence of 25 μg/mL trilostane (TRIL) or of 10 μM DEAB. (*E*-*H*) Retinoid signaling directly stimulates spermatogonial proliferation during spermatogenetic recovery. (*E*) Testis tissue was collected from adult males exposed to 10 nM estradiol (E_2_) *in vivo* for 21 days leading to an interruption of spermatogenesis and accumulation of type A spermatogonia in the context of low Fsh and androgen levels. Testicular explants were then cultured for 2 days in the presence of BMS493 (10 μM) and DEAB (10 μM) inhibitors, followed by another 7 days incubation with different treatments (25 μg/mL TRIL plus 100 ng/mL Fsh) in the absence or presence of 10 μM RE. Testis tissue was collected and used for immunocytochemical detection of BrdU (*F*) and quantification of BrdU labeling indices (*G*), and for quantifying the areas occupied by the type A spermatogonia (*H*). Data are expressed as mean ± SEM (n = 5-12) and asterisks indicate significant differences between groups (* *P* < 0.05, ** *P* < 0.01, *** *P* < 0.001, paired Student’s t test (*A*, *B* and *D*), unpaired Student’s t test (*G* and *H*) or one-way ANOVA followed by Tukey’s multiple comparison test (*C*)). In *D*, results are shown as relative to the basal control condition, which is set at 1 (dashed line). In *F*, scale bar represents 25 μm. A_und_, type A undifferentiated spermatogonia (dashed black line denote positive BrdU cells); A_diff_, type A differentiating spermatogonia (arrowheads indicate positive BrdU cells); SPB, type B spermatogonia; Apop, apoptotic cells. **Figure 8-source data 1.** Raw data used to generate the statistical graphs in Figure 8.

To further investigate retinoid effects on spermatogonial proliferation, we used an experimental model based on estrogen-inhibited Fsh and androgen levels causing an accumulation of type A spermatogonia (36, 59) (see Fig. 8*E*). This testis tissue, enriched for type A spermatogonia, was then used in tissue culture experiments. The cultured testes were first exposed for 2 days to DEAB and BMS493, a pan-RA receptor antagonist, to block both RA synthesis and action. Then, while blocking steroid synthesis, we challenged testis tissue with Fsh and asked if additional RE would modulate Fsh effects. Under these conditions, RE further stimulated the proliferation of type A_und_ and A_diff_ spermatogonia (Figs. 8*F* and *G*), which was mainly differentiating proliferation, as suggested by the decrease of the area occupied by type A_und_ and the increase of the area occupied by type A_diff_ spermatogonia (Fig. 8*H*). These observations suggest that independent of steroid production, Fsh stimulates the differentiating proliferation of type A_und_ and A_diff_ spermatogonia in part via increasing RA synthesis, and moreover that in the presence of Fsh, addition of RE further promotes the differentiation of type A spermatogonia, partially depleting the pool of type A_und_ spermatogonia.

## Discussion

By examining transcriptional changes in the zebrafish testis during initial stages of spermatogenic recovery after pharmaceutical germ cell depletion, we reveal activation of a transcriptional network that includes growth and differentiation factors, together with a stimulation of retinoid and androgen signaling. For the first time in fish, we have identified specific germ cell types responding to RA signaling, finding evidence for a regulatory link between the reproductive hormones Fsh and androgens on one side, and RA production and action on the other (summarized schematically in Fig. 9). Although a *stra8*-like gene is missing in zebrafish, RA regulation is still required for normal spermatogenesis, possibly using other target genes as *rec8a*. Different from mammals, RA signaling in zebrafish is not critical for meiosis, spermatogenesis and sperm production and after an initial period of poor spermatogenesis with age there even is recovery, likely involving androgen-driven compensation. It appears that RA-mediated regulation of testis physiology is conserved in vertebrates, but different mechanisms are used in different taxonomic groups. While RA signaling is strictly required for germ cell survival and development in mammals, fish spermatogenesis still proceeds when RA signaling is disturbed but produces sperm of low quality.

**Fig. 9.**
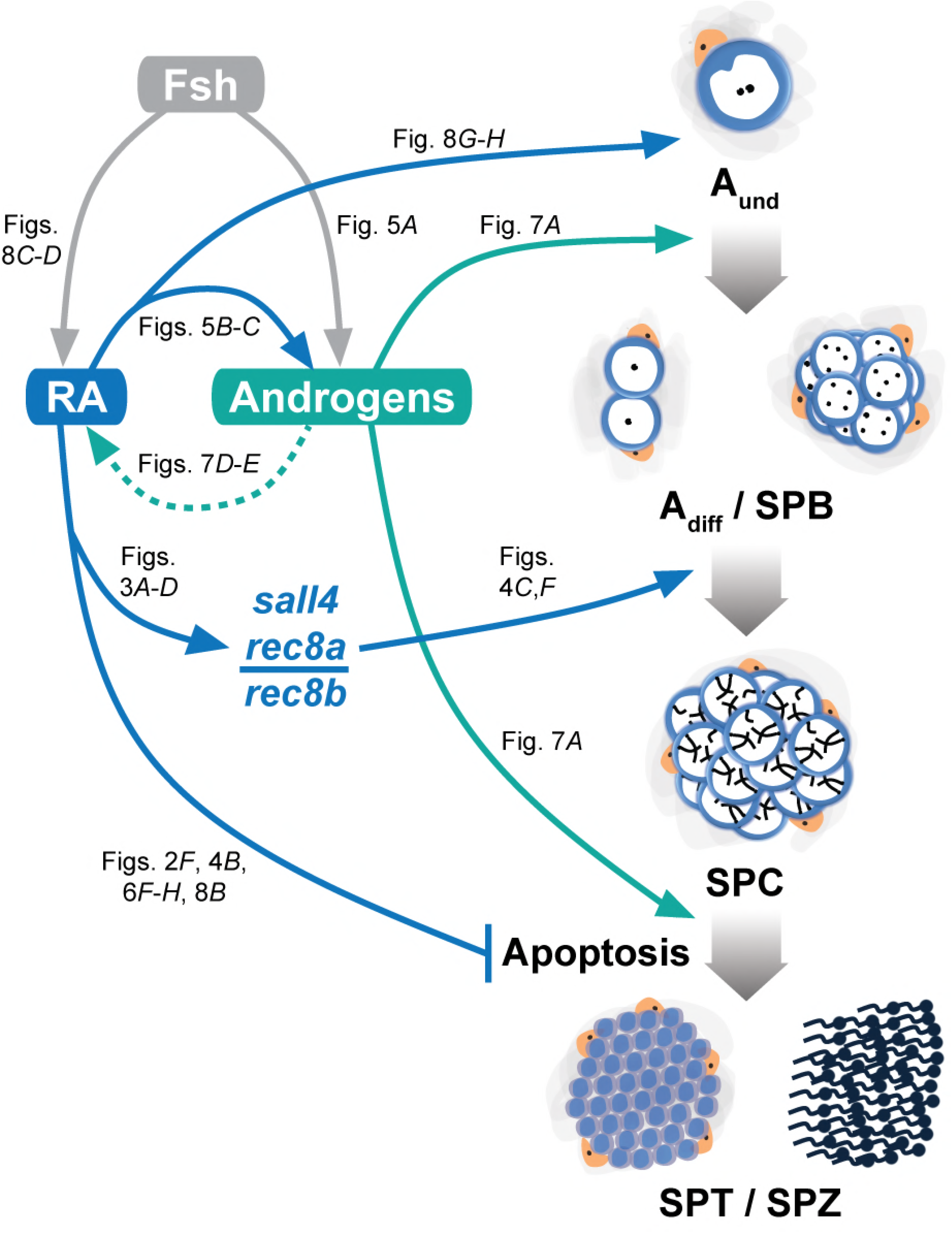
Schematic illustration showing main stages affected by retinoic acid and androgen signaling systems during Fsh-stimulated zebrafish spermatogenesis. Described effects are indicated by arrows (summarizing the results presented in the various figures indicated), and blue (retinoic acid; RA) and green (androgens) lines represent the specific stages influenced. Dashed line denotes assumed effects. Fsh, follicle-stimulating hormone; A_und_, type A undifferentiated spermatogonia; A_diff_, type A differentiating spermatogonia; SPB, type B spermatogonia; SPC, spermatocytes; SPT, spermatids; SPZ, spermatozoa.

### Transcriptomic profiling identifies retinoid and steroid signaling pathway activation during germ cell recovery

Using an established experimental model (54, 60), we created spermatogenic tubules containing Sertoli cells and a small number of A_und_ spermatogonia, probably representing surviving SSCs. We assume that the spermatogenic recovery observed some days later was fueled from these stem cell niches. Depleted testis tissue was characterized by the transcriptional enrichment of the “Retinol metabolism” and of the “Steroid hormone biosynthesis” pathways, the latter being consistent with the enhanced androgen production found in busulfan-depleted testis tissue in our previous study (54). Increased androgen signaling may have supported the spermatogenic recovery, considering the stimulatory effect of androgens on zebrafish spermatogenesis (36, 45, 46) (Fig. 6).

Transcript levels of genes encoding SSC self-renewal factors expressed in Thy^+^ undifferentiated mammalian spermatogonia (55), were up-regulated in depleted zebrafish testes (*Supplementary file 1C*). This gene set includes colony stimulating factor 1 receptors (*csf1ra* and *csf1rb*), while enhanced expression of Csf1 ligand (*csf1a*) was found during recovery (*Dataset 1*). In mice, macrophages, peritubular myoid and Leydig cells in the interstitial tissue adjacent to the seminiferous tubules secrete CSF1 (61), which promotes SSC self-renewal (62) and –in collaboration with RA signaling– differentiation of undifferentiated spermatogonia (63). The morphometric analyses during early recovery showed a remarkable increase in the number of differentiating spermatogonia compared to both busulfan-depleted and untreated control testes. It appears that after SSC regeneration, rapid and widespread differentiation of spermatogonia is the response of the zebrafish testis to the cytotoxic insult, associated with the activation of retinoid and steroid signaling. These coincident changes prompted us to examine experimentally retinoid/androgen signaling interactions.

### Retinoids directly control spermatogonial differentiation in zebrafish in a Stra8-independent manner

Our results indicate that as in mammals (3, 7, 15) in zebrafish too, RA is required for spermatogonial differentiation. However, in zebrafish besides RA also androgens are crucial for spermatogonia to differentiate, as A_und_ spermatogonia accumulate when androgen production is blocked. This arrest of spermatogonial differentiation in the absence of androgens occurs despite the presence of RE, indicating that both retinoids and androgens are needed to induce differentiation. Besides its role in the induction of spermatogonial differentiation, RE also stimulates proliferation of type A_und_ and A_diff_ spermatogonia in testis tissue enriched for type A spermatogonia after administration of Fsh. This effect was androgen-independent as it could not be blocked by inhibition of steroid production with trilostane (Figs. 8*E*-*G*). Under similar circumstances (i.e. using testis tissue enriched in type A spermatogonia), spermatogonial differentiation was promoted by Fsh-stimulated release of Insl3 and Igf3 (49). In the present context, it seems possible that Fsh creates a growth factor environment, in which exogenous RE not only further stimulates the proliferation of type A_diff_, but also of type A_und_ spermatogonia, promoting differentiating proliferation in both type A spermatogonia.

The mechanism by which RA stimulates spermatogonial differentiation in mice involves KIT (reviewed in (2)). This may be different in zebrafish, where we did not observe a clear modulation of Kit ligand transcript levels (*Supplementary file 6*). Moreover, RA action also did not involve changes in transcript levels of growth factors known to modulate germ cell differentiation in zebrafish (e.g. Igf3 (51, 64), Insl3 (48, 49) and Amh (53, 64, 65)). Instead, we found that the previously described mouse RA-regulated germ cell factors, SALL4A (5, 66) and REC8 (6), may mediate induction of differentiation in the zebrafish testis. Our gene expression studies on FACS-enriched zebrafish germ cells suggest direct retinoid effects on transcript levels of these genes in germ cells. In mice, the germ cell-specific expression of *Sall4a* is controlled by RARγ (5) and its genetic ablation in spermatogonia results in the loss of differentiating germ cells (66). In zebrafish, RA signaling also increased *sall4* transcript levels in spermatogonia, and knockdown of *sall4* decreased expression of germ cell-marker (and steroidogenesis-related) genes, which is in line with reduction of differentiating spermatogonial cell numbers after blocking RA synthesis. REC8 is a meiosis-specific cohesin component expressed in early spermatocytes in rodents (67, 68). Loss of the *Rec8* locus in mice results in high mortality rates and sterility in both sexes; in males surviving until adulthood, haploid germ cells are absent (69). Since both isoforms of the *rec8* gene are up-regulated by retinoids, it is possible that Rec8 mediates RA-stimulated germ cell development also in zebrafish. Indeed, knockdown of *rec8a* and *rec8b* causes a significant decrease on germ cell-marker gene expression, suggesting that *rec8* may mediate part of the RA effects in zebrafish germ cells.

### Germ cell-specific inhibition of RA signaling

In our dominant-negative transgenic zebrafish model with a germ cell-specific inactivation of RA signaling, at 6 months of age there is partial germ cell depletion and increased apoptosis. Still, spermatogonial differentiation, meiosis and the production of mature sperm occurs, resulting in a quite heterogeneous testis histology. This differs from the generalized block of spermatogonial differentiation found in mice using a similar transgenic approach (15). However, sperm quality was clearly compromised in these transgenic zebrafish, with reduced fertilization rates and a high mortality of the (few) resulting embryos within the first 24 hours of development. Unexpectedly, at 9 months of age spermatogenesis recovers in terms of testis histology, and a testis weight clearly above wild type control levels. The latter seems related to the prominent increase in the numbers of interstitial cells in transgenic zebrafish. We speculate that the accompanying enhanced levels of steroidogenesis-related genes and androgen receptor transcripts, indicate that Leydig cell activation occurs, allowing an androgen-mediated spermatogenic recovery. However, the spermatogenic recovery in transgenic zebrafish did not improve the low sperm quality. More work is required to clarify what signaling mechanisms are involved in this response and their significance for testis physiology. Considering the three candidate genes studied by GapmeR-mediated knockdown, we assume that in particular *rec8a* plays an important role in the germ cells’ response to retinoids, as indicated by its consistent down-regulation in the testes of *dn-raraa* transgenic fish. Interestingly, knockdown of *rec8a* (but not *rec8b*) also led to a significant decrease of steroidogenesis-related transcripts, which may be related to the reduced testicular androgen release after blocking RA production (Fig. 5). Taken together, our functional evidence shows that the germ cell-specific lack of RA signaling results in a clear spermatogenesis phenotype in young adults that partially recovers within 3 months, associated with Leydig cell over-activation, and that Rec8 may be one of the mediators of RA action in the zebrafish testis.

### Androgens and retinoids jointly promote spermatogenesis

Next to retinoids, also androgens stimulate vertebrate spermatogenesis (29, 70). In mice, Sertoli cell-selective knockout of the androgen receptor (SCARKO) blocks meiosis (28), while its ubiquitous ablation (ARKO) reduces the numbers of spermatogonia in adulthood (71). Also zebrafish Sertoli cells express the androgen receptor (72), and 11-KT (which is the main androgen in fish) initiated spermatogonial differentiation in tissue culture or during puberty in other fish species (33, 34). In zebrafish deficient for the androgen receptor adult males show clearly smaller testes than WT fish, though still some spermatozoa are produced in these fish(45, 46). In contrast, in mice meiosis and spermiogenesis strictly depend on androgen signaling (28, 29, 70). The present study shows that androgen-induced increases in transcript levels of mitotic and meiotic germ cell-markers are compromised by blocking RA production, which also elevates apoptosis among spermatocytes and in particular spermatids (Fig. 6). Blocking steroid production in the presence of RE, on the other hand, leads to an accumulation of type A_und_ spermatogonia and spermatocytes (Fig. 7). This suggests that retinoid-stimulated production of type A spermatogonia and meiotic cells is complemented by androgen signaling that promotes spermatogonial differentiation and completion of meiosis, respectively. Therefore, we propose that androgens and retinoids target developmental steps during spermatogenesis in a complementary manner to guarantee efficient sperm production in zebrafish. Such an androgen/retinoid interplay in regulating the differentiation of spermatogonia is a new observation in the vertebrate testis. However, interactions between androgen and RA signaling were demonstrated in the lacrimal gland (73), hippocampus (74) and prostate carcinoma cells (75). Different from mammalian models, where preventing either androgen or retinoid signaling completely blocked spermatogenesis, removing one of these pathways in zebrafish still allows (some) sperm production to occur, albeit in a compromised manner.

In zebrafish, an important effect of blocking retinoid signaling is to increase apoptotis of spermatocytes and in particular spermatids, while there is no block in spermatogonial differentiation and entry into meiosis of spermatocytes. In contrast, in mammals, spermatogonial differentiation and entry into meiosis are completely blocked in the absence of RA signaling, and a potential interaction of RA signaling on androgen action has not been demonstrated so far. The meiotic block in ARKO mice is associated with increased germ cell apoptosis (76), but this seems to be a phenomenon independent of RA signaling. In zebrafish, inhibition of steroid production in the presence of RE has no effect on germ cell apoptosis, while the androgen-induced decrease of apoptosis (Fig. 6*C*) may reflect androgen support of RA production. Thus, our data suggest that 11-KT and RA promote the production of differentiating spermatogonia, while 11-KT (but not RA) also promotes the completion of meiosis, and RA (but not 11-KT) increases the efficiency of spermiogenesis by reducing the apoptotic loss of spermatids.

### Fsh increases RA production and uses retinoid signaling to stimulate spermatogenesis

Fsh is an important hormone regulating spermatogenesis also in zebrafish (49). Its action in the fish testis is mediated by Sertoli and Leydig cells, both expressing the *fshr* gene (47, 77-79), so that Fsh is a potent steroidogenic hormone (47, 78, 80, 81) and also modulates growth factor production (49). We now have shown that another role of Fsh is to increase testicular RA production, thereby stimulating RA target gene transcript levels. Hence, the biological activity of Fsh in the zebrafish testis may, in part, be mediated by RA signaling. *Vice versa*, RA biosynthesis seems to benefit from Fsh-stimulated androgen production considering that *aldh1a2* (encoding the RA-producing enzyme in zebrafish (17, 82)) transcript levels increase upon androgen treatment under *ex vivo* and *in vivo* conditions. The also concomitant androgen-mediated increases of *cyp26a1* suggest that complex mechanisms for RA production and degradation operate in the adult testis, similar to other tissues in zebrafish (83).

## Material and Methods

### Animals

Sexually mature wild type (WT; AB strain) and transgenic *Tg(vasa:EGFP)* (84) and *Tg(vasa:raraa_S392stop-IRES-mCherry, myl7:EGFP)* –referred to as *dn-raraa*– zebrafish (*Danio rerio*) males, were used for the experiments described in the present study. For the construction of the transgenic line *dn-raraa*, the amino acid sequence of the endogenous zebrafish Raraa (the most similar isoform to human RARα; ~80% similarity) was aligned with the amino acid sequence of a dominant-negative mutant form of the human RARα previously described (RARα403*; (56)). This truncated receptor (Raraa^DN391^) protein misses the C-terminal domain required for interacting with cofactors to activate transcription, but is able to dimerize with endogenous Rar and Rxr proteins and then bind to RARE (RA response element) motifs. First, the *raraa^DN391^* cDNA sequence was PCR-amplified (Advantage 2 PCR Kit, Clontech) using testis cDNA and subcloned into pME-MCS (85). Using Gateway technology, the *raraa^DN391^* sequence, followed by an internal ribosome entry sequence (IRES) and the mCherry-coding region, was placed under the control of the *vasa* promoter and inserted into pDestTol2CG2 (85). The latter plasmid includes a cardiac myosin light chain 7 regulatory (*myl7*) enhancer-promoter gene driving EGFP that can be used as screening tool.

Handling and experimentation were consistent with the Dutch national regulations and the Life Science Faculties Committee for Animal Care and Use in Utrecht (The Netherlands) approved the experimental protocols.

### Transcriptomic analysis by RNAseq

Testicular samples considered for RNAseq studies were obtained employing the experimental model shown in *Supplementary file 1A*. Upon an acclimatization period with increasing temperature (from 27 to 35°C; ~1°C increment/day), adult zebrafish males were exposed to 35°C for 14 days and injected with the cytostatic agent busulfan (single intraperitoneal injection after 7 days at 35°C; 40 mg/Kg). Then, fish were placed back to normal water temperature and testis samples were collected at different time points. Morphological analysis of testicular samples showed maximum germ cell depletion 10 days post busulfan injection (i.e. 10 dpi; *Supplementary file 1Biii*) and the recovery of endogenous spermatogenesis ~14 dpi (*Supplementary file 1Biv*).

Total RNA was isolated from 1) testes of untreated adult control zebrafish, 2) germ cell-depleted, and 3) testis tissue at the beginning of the recovery period, using the miRNeasy Mini Kit (Qiagen) according to the manufacturer’s protocol. RNA integrity was checked with an Agilent Bioanalyzer 2100 total RNA Nano series II chip (Agilent). Only samples with a RNA integrity number > 8 were used for library preparation. Illumina RNAseq libraries were prepared from 2 μg total RNA using the Illumina TruSeq RNA Sample Prep Kit v2 (Illumina, Inc.) according to the manufacturer’s instructions. The resulting RNAseq libraries were sequenced on an Illumina HiSeq2500 sequencer (Illumina, Inc.) as 1×50 nucleotide reads. Image analysis and base calling were done by the Illumina pipeline. Quality control of the obtained reads was performed using FastQC suite (v0.10.1; default parameters). The sequencing yield ranged between ~11 and ~70 million reads per sample and mapping efficiency for uniquely mapped reads was between 64.8 and 73.3% (see *Dataset 1*). RNAseq derived reads were aligned to the zebrafish genome (Zv9) using TopHat (v2.0.5; (86)). The resulting files were filtered with SAMtools (v0.1.18; (87)), and the read counts extracted using the Python package HTSeq (www-huber.embl.de/HTSeq/doc/overview.html/; (88)). Data analysis was performed with the R/Bioconductor package DESeq (P < 0.005, fold change [FC] ≥ 3.0; (89)). The raw RNAseq data of the 15 samples sequenced (5 biological replicates per condition) have been deposited in the NCBI GEO database with accession number GSE116611. Regulated KEGG pathways were determined using the KEGG Mapper tool (90). KEGG pathways represented by at least 5 differentially expressed genes (DEGs) and by the ratios of regulated genes (up-/down-, and *vice versa*) higher than 5 were considered for the analysis.

Functional enrichment analyses were carried out using a plugin available at http://www.baderlab.org/Software/EnrichmentMap/ (91) for the Cytoscape network environment (92). The Enrichment Map plugin calculates over-representation of genes involved in closely related Gene Ontology (GO) categories (93), resulting in a network composed of gene sets grouped according to their function. DAVID Bioinformatics Resources 6.7 (http://david.ncifcrf.gov/; (94)) was used to retrieve GO terms from the list of DEGs and exported as the input for each functional enrichment analysis.

### Testis tissue culture

Using a previously established *ex vivo* culture system (37), adult zebrafish testis tissue was incubated in the presence of various test compounds to investigate retinoid involvement in spermatogenesis including retinoic acid (RA), retinol (RE, RA precursor), 4-diethylaminobenzaldehyde (DEAB, RA production inhibitor (95, 96)) and BMS493 (pan-RA receptor antagonist (97, 98)). All these compounds were purchased from Sigma-Aldrich and used at the same final concentration of 10 μM. Additional *ex vivo* experiments were performed to test the effects of recombinant zebrafish Fsh (100 ng/mL (47)) and 11-ketotestosterone (11-KT, 200 nM; Sigma-Aldrich). When focusing on the effects not mediated by steroid hormones, incubations were carried out in the presence of trilostane (TRIL, 25 μg/mL; Sigma-Aldrich), which prevents the production of biologically active steroids. Moreover, an androgen insufficiency model was employed to obtain the results presented in Figs. 8*E*-*G*. Briefly, before collecting testis tissue for *ex vivo* culture, the fish were exposed to 10 nM 17-β estradiol (E2; Sigma-Aldrich) *in vivo* for 3 weeks with a daily change of the E_2_-containing water. Using this approach, endogenous spermatogenesis is interrupted such that type A spermatogonia accumulate while the testes become depleted of type B spermatogonia, spermatocytes and spermatids (59).

### Testicular cell suspension and culture

To investigate the cellular localization of testicular *sall4*, *rec8a* and *rec8b* expression, we used testes from transgenic *Tg(vasa:EGFP)* zebrafish expressing enhanced green fluorescent protein under the control of the germ cell-specific *vasa* promoter (84). Testis tissue from ~10 transgenic fish was digested with 1 mg/mL collagenase/dispase solution at 27°C for 2 hours with gentle shaking (99). The resulting cell suspension was filtered through a 70 μm filter and subsequently through a 40 μm filter (BD Bioscience) before centrifugation at 500 rpm for 10 min. The cell pellet was resuspended in PBS (phosphate-buffered saline; pH 7.4) and the resulting suspension immediately submitted to fluorescence-activated cell sorting (FACS) using an in Flux cell sorter (BD Bioscience). Autofluorescence was removed through the FACS dot plot profile generated with a testicular cell suspension from WT males. EGFP-positive and -negative cells were collected, centrifuged in PBS at 1000 rpm for 10 min and the pellet stored at −80°C until use.

In addition, testicular cell suspensions were performed using WT testes and then submitted to culture conditions. Briefly, ~5×10^5^ cells/mL were transferred to 25 mL culture flasks and cultured in L-15 supplemented medium (Gibco) containing 2% UltroserTM G serum substitute (Pall Corporation) at 27°C for 3 days. Using this method, somatic cells adhere to the bottom of the plate while germ cells remain in suspension. Subsequently, an equal number of cells (including both germ and somatic cells after a brief trypsin wash) were transferred to 12 well plates and cultured at 27°C for 3 days in basal, RA- (10 μM) and RE-treated (10 μM) conditions. Upon incubation, cells were centrifuged at 1000 rpm for 10 min and the pellet stored at −80°C until use.

### *In vivo* exposure to androgen

Adult male fish were submitted to water containing 100 nM 11-ketoandrostenedione (OA) or control untreated conditions *in vivo* for 5 weeks with a daily change of water (as described in (36)). Stock solutions were prepared in deionized water by extensive stirring at 37°C, which were further diluted to the proper concentration in aquarium water. Upon *in vivo* treatment, testis tissue was collected and used for gene expression analysis.

### Sample preparation and analysis

Under some of the experimental conditions described in this study (Testis tissue culture section), type A and type B spermatogonia proliferation activity was investigated by studying the incorporation of the S-phase marker bromodeoxyuridine (BrdU; 50 μg/mL, Sigma-Aldrich), which was added to the medium during the last 6 hours of the culture period. After incubation, testis tissue was fixed at room temperature for 1 hour in freshly prepared methacarn (60% [v/v] methanol, 30% chloroform and 10% acetic acid glacial; Merck Millipore) and processed for subsequent analysis. To quantify spermatogonial proliferation, the mitotic index was determined by examining at least 100 germ cells/cysts, differentiating between BrdU-labeled and unlabeled cells. To evaluate the proportion of area occupied by different germ cell types in 4% glutaraldehyde fixed tissue (4°C, overnight), 10-15 randomly chosen fields were photographed at ×400 magnification and the images were analyzed quantitatively using ImageJ software. With a specific plugin, a 540-point grid was made to quantify the proportion of the area for the various germ cell types, based on the number of points counted over those germ cell types. For both purposes (BrdU incorporation and area analyses), testis tissue was dehydrated, embedded in Technovit 7100 (Heraeus Kulzer), sectioned at a thickness of 4 μm. The germ cells/cysts were identified according to previously published morphological criteria (100).

Additional *ex vivo* experiments and testicular cell suspensions were carried out to investigate candidate gene expression in response to different experimental conditions. For all these experiments, total RNA was isolated using the RNAqueous Kit (Ambion) following the manufacturer’s instructions. Relative mRNA levels of candidate genes were analyzed by real-time, quantitative PCR (qPCR; see *Supplementary file 7* for detailed primer information) as previously described (47). The geometric mean of *eef1a1l1*, *rpl13a* and *ubc* was used as housekeeping endogenous control due to their constant expression under the conditions analyzed.

Furthermore, incubation medium was collected to quantify testicular androgen (11-KT) and RA release by radioimmunoassay (as described in (47)) and enzyme immunoassay (commercial kit, MyBioSource), respectively.

### Apoptosis detection by TUNEL

To determine the incidence of apoptosis, paraffin embedded testis tissue that was previously incubated in the absence or presence of the test compounds was subjected to deoxynucleotidyl transferase-mediated dUTP nick-end labeling (TUNEL). First, testis tissue was washed with PBS and subsequently fixed in 4% PBS-buffered paraformaldehyde (4°C, overnight). After a 30 min wash with PBS, testis tissue was dehydrated and embedded in paraffin. Sections of 4 μm thickness were treated with permeabilization solution (0.1% Triton X-100, 0.1% sodium citrate) for 8 min. Finally, testis tissue was incubated with TUNEL reaction mixture (In Situ Cell Death Detection Kit, Fluorescein; Roche) in the dark at 37°C for 1 hour. After washing twice in PBS, sections were counterstained with DAPI, mounted in Vectashield antifade mounting medium (Vector Laboratories) and analyzed by confocal laser scanning microscopy (Zeiss LSM 700). Negative and positive controls were included in each experimental set up (*Supplementary file 5*).

### *sall4*, *rec8a* and *rec8b* knockdown by GapmeR technology

*sall4*, *rec8a* and *rec8b* gene knockdown was investigated by using specific antisense oligonucleotides (LNA™ GapmeRs; Exiqon). GapmeRs were dissolved in the incubation medium at 1 μM final concentration and applied to *ex vivo* tissue culture conditions for 4 days. Unassisted GapmeR uptake method, called gymnosis, has been previously described (101, 102). Three technical replicates per gene were performed and knockdown efficiency was confirmed by qPCR analysis, and normalized to the negative GapmeR control.

### Statiscal analysis

GraphPad Prism 5.0 package (GraphPad Software, Inc.) was used for statistical analysis. Significant differences between groups were identified using Student’s t test (paired or unpaired, as appropriate) or one-way ANOVA followed by Tukey’s test for multiple group comparisons (*, *P* < 0.05; **, *P* < 0.01; ***, *P* < 0.001; ns, no significant changes observed). Data are represented as mean ± SEM.

## Acknowledgements

The authors thank Rafael H. Nóbrega (Institute of Bioscience of Botucatu, São Paulo, Brazil) for assistance during busulfan *in vivo* experiments, Daniëlle Janssen (Utrecht University aquarium facility) for maintaining the zebrafish stocks and for technical support and Ger J. A. Arkesteijn (Faculty of Veterinary Medicine, Utrecht, The Netherlands) for expert assistance with the FACS analysis. This study was co-funded by the Research Council of Norway BIOTEK2021/HAVBRUK program with the projects SALMAT (n° 226221) and SALMOSTERILE (n° 221648).

## Supplementary file legends

**Supplementary file 1.** Germ cell-depleted testes rapidly recover following a cytotoxic insult. (*A*) Schematic representation of the experimental model inducing germ cell depletion/recovery of spermatogenesis. Following an acclimatization period with increasing temperature (from 27 to 35°C; ~1°C increment/day), adult zebrafish males were exposed to 35°C for 14 days and injected with the cytostatic agent busulfan (single intraperitoneal injection after 7 days at 35°C; 40 mg/Kg). Then, fish were placed back to normal water temperature and testis samples collected at different time points. (*B*) Morphological analysis showed maximum germ cell depletion 10 days post busulfan injection (i.e. 10 dpi; panel *iii*) and ongoing recovery of spermatogenesis ~14 dpi (panel *iv*; scale bar represents 25 μm). Inset *iii* shows a single type A undifferentiated (A_und_) spermatogonium (white arrowhead; scale bar represents 10 μm). (*C*) Germ cell-marker expression in depleted *versus* recovering testes. mRNA abundance of undifferentiated (55) and differentiating germ cell-marker genes identified by RNAseq and retrieved in the different analyses used in this study (for each comparison, the control condition appears in brackets). Genes differentially expressed (n = 5; *P* < 0.005, [FC] ≥ 3.0; see Dataset 1 for detailed information) are highlighted with red or green (up- or down-regulation, respectively) background. A_und_, type A undifferentiated spermatogonia; A_diff_, type A differentiating spermatogonia; SPB, type B spermatogonia; SPC, spermatocytes; SPT, spermatids.

**Supplementary file 2.** Transcriptomic analyses of zebrafish testes before (Control) and at different times (10 days, Depletion; 14 days, Recovery) after a cytostatic insult. Total numbers of up (red)- and down (green)-regulated genes, and KEGG pathways, identified by RNAseq during depletion (*A*) and subsequent recovery (*B*) (n = 5, *P* < 0.005, fold change [FC] ≥ 3.0; see Dataset 1 for detailed information). Each KEGG pathway shown is represented by at least 5 differentially expressed genes (DEGs) and has a ratio of (up-/down-, or vice versa) regulated genes higher than 5. DEGs are highlighted with red (up-) or green (down-regulated) background.

**Supplementary file 3.** Testicular cell suspensions. (*A*) Retinoid effects on germ cell (Germ)- and retinoic acid (RA)-related gene expression in cell culture conditions. Testicular cell suspensions were obtained as previously described (99) using wild type (WT) tissue from adult males (20 testes) and then submitted to culture conditions. Cells were cultured in L-15 supplemented medium containing 2% UltroserTM G serum substitute for 3 days at 27°C. Subsequently, an equal number of cells (1×10^5^/mL; including both germ and somatic cells after a brief trypsin wash) were cultured for 3 additional days at 27°C in basal medium, or in medium containing 10 μM RE or RA. After incubation, cells were collected (brief trypsin wash) and RNA was isolated for qPCR analysis. Data are shown as mean of fold change ± SEM (n = 3; technical replicates) and expressed relative to the basal condition, which is set at 1. Asterisks indicate significant differences between treated and control groups (* *P* < 0.05, ** *P* < 0.01, *** *P* < 0.001, unpaired Student’s t test). (*B*) qPCR analysis of EGFP- and EGFP+ cell fractions. A pooled testicular suspension was obtained from transgenic *Tg(vasa:EGFP)* testes, expressing enhanced green fluorescent protein under the control of the germ cell-specific *vasa* promoter (84), and subjected to FACS. Dot plot shows the total suspension from which a population of large cells (x-axis) showing an intense fluorescence (y-axis) was harvested (54), as indicated by the shape delineated with a black line (EGFP^+^ cell fraction). Autofluorescence was removed through the FACS dot plot profile generated with a testicular cell suspension from WT males. EGFP^-^ and EGFP^+^ cell fractions were collected and RNA was isolated for qPCR analysis of germ and somatic cell markers, as well as RA-related genes. Data are shown as mean fold change ± SEM (n = 3; technical replicates) and expressed relative to the EGFP^-^ condition, which is set at 1 (* *P* < 0.05, *** *P* < 0.001, unpaired Student’s t test).

**Supplementary File 3-source data 1.** Raw data used to generate the statistical graphs in Supplementary File 3.

**Supplementary file 4.** Abnormal spermatogenesis in *dn-raraa* transgenic zebrafish testes. (*A*) Confirmation of transgenesis in adult *dn-raraa* males (microinjected as 1 cell stage embryo). Representative *myl7*-driven green fluorescent (EGFP-positive) heart and *vasa*-driven red fluorescent (mCherry-positive) testis tissue are shown and compared with WT sibling tissues. Representative PCRs showing the absence or presence of the expected band (indicated by an arrowhead) for the *raraa^DN391^* transgene in samples generated from WT and *dn-raraa* adult males, respectively. (*B*) Heterogeneity of testis histology in 6 month-old *dn-raraa* transgenic males. For each individual (n = 4), two different areas of the seminiferous epithelium −at two different magnifications− are shown. In panels at 200X magnification, white asterisks denote germ cell-depleted tissue while black asterisks indicate empty areas. Scale bar represents 25 μm.

**Supplementary file 5.** Inhibition of retinoic acid (RA) production in the presence of androgen increases germ cell apoptosis. (*A*-*I*) Representative images of testis tissue subjected to TUNEL analysis. Incidence of apoptosis in paraffin sections from testicular explants cultured for 4 days in the presence of 200 nM 11-KT and 10 μM DEAB (the respective control –group treated only with 11-KT− is shown in Fig. 6*H*). Negative (TUNEL reaction mixture without enzyme solution; *D*-*F*) and positive (recombinant DNase I; *G*-*I*) controls were developed according to the manufacturer’s indications. Green staining indicates fragmented DNA in cells undergoing apoptosis (TUNEL+) and blue staining indicates DNA (DAPI counterstain). Insets in *A*-*C* show a representative apoptotic cyst with higher magnification.

**Supplementary file 6.** Retinoid action in zebrafish testis is not mediated by the Kit signaling pathway. (*A* and *B*) Changes in testicular transcript levels of Kit ligands (*kitlga* and *kitlgb*) and receptors (*kita* and *kitb*) in response to 10 μM RE (*A*) or after blocking RA production by adding 10 μM DEAB (*B*). (*C* and *D*) Kit-related gene expression was also studied in response to 11-KT compared to basal conditions (*C*), and to 11-KT in the absence or presence of 10 μM DEAB (*D*). Data are shown as mean of fold change ± SEM (n = 6-12) and expressed relative to the control condition, which is set at 1. Asterisks indicate significant differences between groups (* *P* < 0.05, paired Student’s t test).

**Supplementary File 6-source data 1.** Raw data used to generate the statistical graphs in Supplementary File 6.

**Supplementary file 7.**
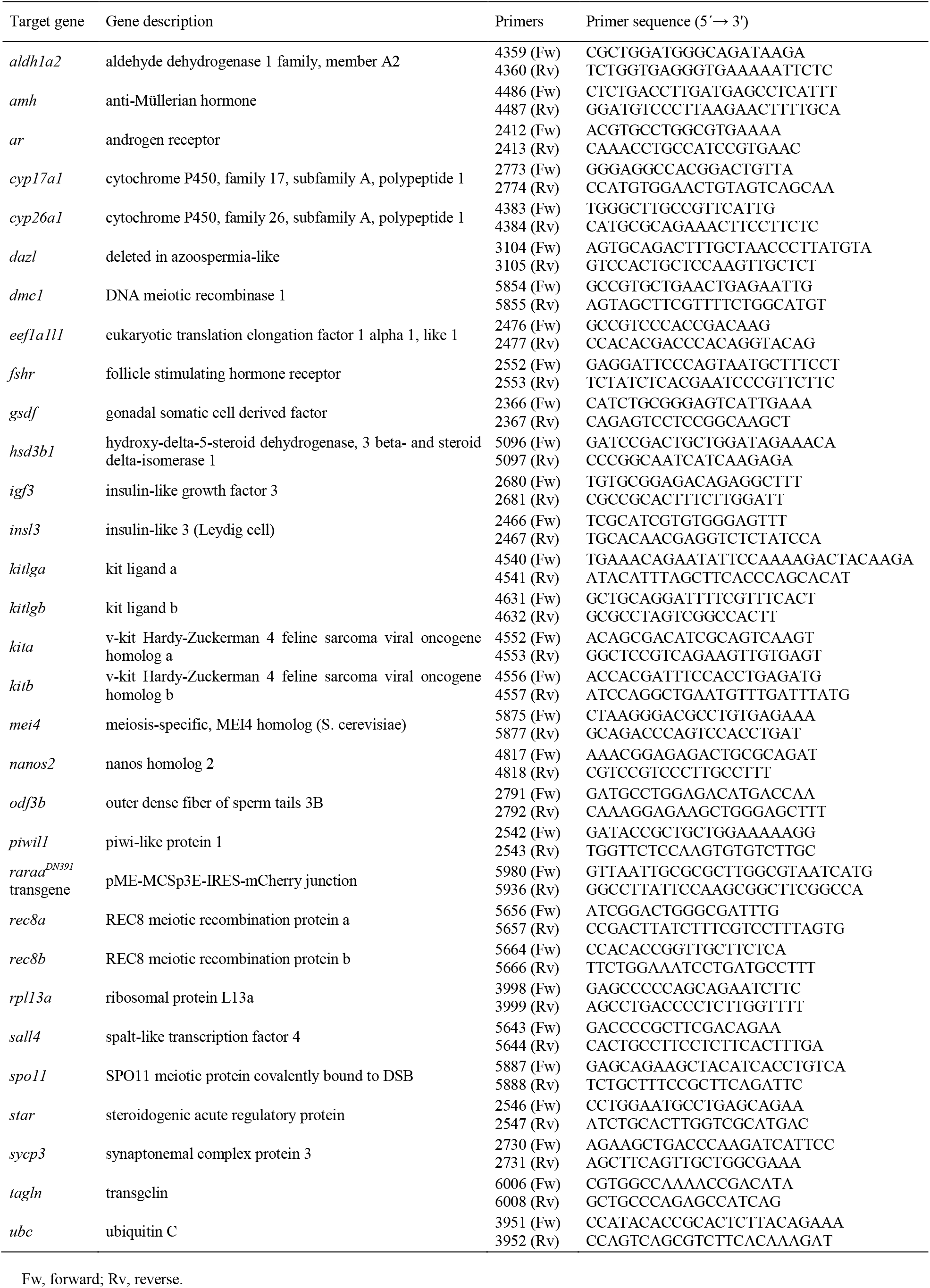
Sequences of primers used in gene expression analyses by qPCR.

## Additional supporting data files

**Dataset 1.** Characterization of RNAseq data. Summary of read counts and mapping statistics for each RNAseq replicate. The file also contains complete sets of modulated genes identified (n = 5, *P* < 0.005, fold change [FC] ≥ 3.0, unpaired Student’s t test) in the zebrafish testis in the different comparisons considered in this study.

